# Constructing Biologically Constrained RNNs via Dale’s Backprop and Topologically-Informed Pruning

**DOI:** 10.1101/2025.01.09.632231

**Authors:** Aishwarya H. Balwani, Alex Q. Wang, Farzaneh Najafi, Hannah Choi

## Abstract

Recurrent neural networks (RNNs) have emerged as a prominent tool for modeling cortical function, and yet their conventional architecture is lacking in physiological and anatomical fidelity. In particular, these models often fail to incorporate two crucial biological constraints: i) Dale’s law, i.e., sign constraints that preserve the “type” of projections from individual neurons, and ii) Structured connectivity motifs, i.e., highly sparse yet defined connections amongst various neuronal populations. Both constraints are known to impair learning performance in artificial neural networks, especially when trained to perform complicated tasks; but as modern experimental methodologies allow us to record from diverse neuronal populations spanning multiple brain regions, using RNN models to study neuronal interactions without incorporating these fundamental biological properties raises questions regarding the validity of the insights gleaned from them. To address these concerns, our work develops methods that let us train RNNs which respect Dale’s law whilst simultaneously maintaining a specific sparse connectivity pattern across the entire network. We provide mathematical grounding and guarantees for our approaches incorporating both types of constraints, and show empirically that our models match the performance of RNNs trained without any constraints. Finally, we demonstrate the utility of our methods for inferring multi-regional interactions by training RNN models of the cortical network to reconstruct 2-photon calcium imaging data during visual behaviour in mice, whilst enforcing data-driven, cell-type specific connectivity constraints between various neuronal populations spread across multiple cortical layers and brain areas. In doing so, we find that the interactions inferred by our model corroborate experimental findings in agreement with the theory of predictive coding, thus validating the applicability of our methods.

## 1 Introduction

Recent years have seen the increasing adoption of artificial neural networks (ANNs) for modeling brain function both mechanistically and algorithmically [1, 2, 3, 4, 5]. In particular, recurrent neural networks (RNNs) are now an established tool in computational neuroscience research [6, 7], being used to study neuronal computation at varying scales ranging from subsets of neurons sampled across a single brain region, two interacting regions [8, 9, 10], and even numerous populations spread across multiple interacting brain regions [11, 12]. By way of either reproducing desired behaviours [13, 14, 15], task-driven responses [16, 17, 18], or by fitting to recorded neural data [19, 20], RNNs have been shown to successfully capture latent dynamics typical of neural circuits [21, 22] thus making them especially useful for modeling phenomena observed across the cortex. The degree to which ANNs can effectively approximate neural data however depends on two key considerations: i) The literature suggests a direct correlation between the ability of an ANN to learn well on a task and the extent to which its behaviour and learnt representations match real neural data [23, 24, 25], and ii) More biologically realistic architectures aid in the learning of representations that better match real neuronal data [26, 27, 28, 29]. These factors make it essential that the ability of the ANNs to learn and represent a wide range of function classes be unrestricted, that their training not suffer from hindrances, and their construction respect important anatomical principles, especially when being used as models of the brain to study neuroscientific phenomena [30, 31].

Of the various discrepancies conventional RNN-based neuroscientific models have with their biological counterparts (Fig. 1.A1) [32], two notable ones are their lack of adherence with Dale’s principle [33], i.e., the phenomena that restricts a presynaptic neuron to have exclusively either an excitatory or inhibitory effect on all its postsynaptic connections, and structured sparse connectivity amongst neuronal populations, a fundamental feature of brain organization observed across various species [34, 35, 36] and brain regions [37]. Unfortunately, directly incorporating these constraints oftentimes decreases the capacity and flexibility of the network to fit the training data, which leads to a drop in learning performance [38, 39]. While there has been active research towards addressing these issues in both the machine learning (ML) [40, 41, 42, 43] and computational neuroscience communities [38, 44, 45, 46, 47, 48, 49], these efforts have mostly been made to include these constraints into the network structure individually, rather than in conjunction as one would see in biological brains [50]. Subsequently, there remains a need for ways to construct sparse, sign-constrained deep neural networks which can also achieve performance levels comparable to conventional ANNs.

**Figure 1:**
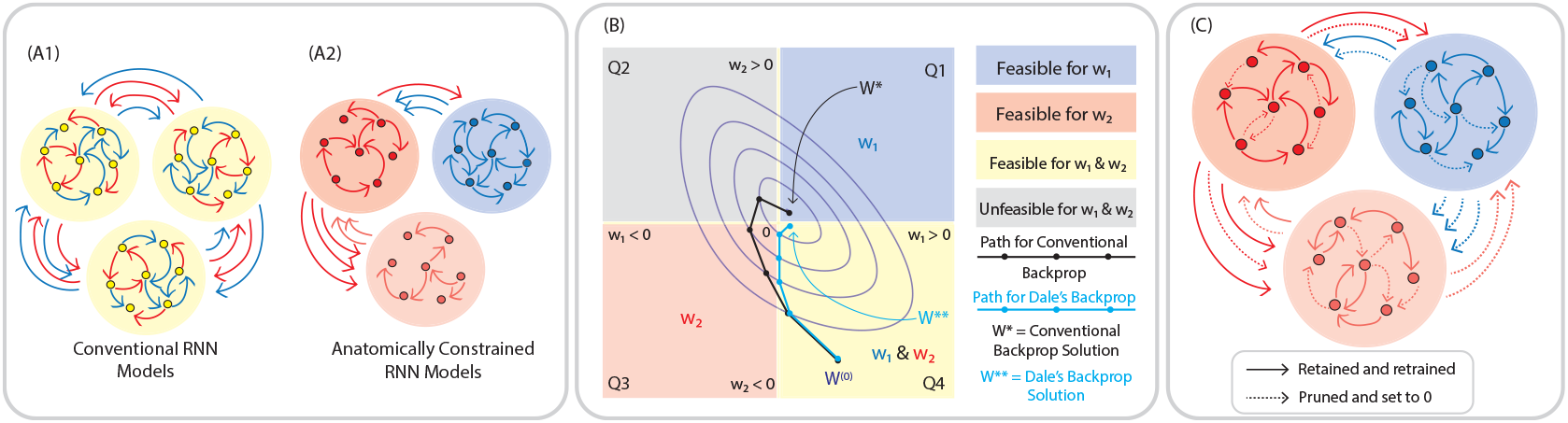
Schematic for Constructing Biologically Constrained RNN Models. **(A)** Illustration of conventional vs. biologically constrained RNN models (A1 vs. A2). Conventional RNNs consist of general purpose neurons that project a mix of excitatory and inhibitory signals, with no specific connectivity structure within or across populations. Biologically constrained RNNs restrict populations of neurons to be either strictly excitatory (red) or inhibitory (blue), with anatomically-informed connectivity motifs both within and across populations. **(B)** Optimization in parameter space when training with conventional backpropagation (black) vs. Dale’s backprop (blue). Purple contours represent level sets for the positions the algorithms take in parameter space at different time steps. **(C)** Enforcing anatomically-consistent connectivity motifs. Dashed lines represent connections that are set to 0 during the pruning process. Solid lines represent connections that are retained post-pruning.

Our work therefore introduces methods that allow us to easily incorporate both neuronal sign constraints and sparse connectivity motifs into the conventional backpropagation-based RNN training pipeline (Fig. 1.A2). Specifically, we first train a dense network that respects a pre-determined set of sign constraints via a modified version of standard backpropagation [51] which we call *Dale’s backpropagation* (Fig. 1.B), after which we prune away weights using a probabilistic pruning rule we call *top-prob pruning* (Fig. 1.C), to achieve a target connectivity pattern. Finally, we retrain the sparse sub-network retained post pruning once again with Dale’s backprop. Importantly, both of our methods are mathematically grounded and follow rigorous principles. With Dale’s backprop, we provide theoretical guarantees on the linear convergence of the algorithm under specific conditions, ensuring that the training respects anatomical constraints while achieving optimal learning performance. Our pruning rule is motivated by topological principles, particularly the preservation of high-magnitude weights that contribute to the network’s zeroth-order connectivity structure, thus enhancing the functional and anatomical plausibility of the model.

Besides being convenient, easily implementable, and scalable using standard ML packages, our approach also aligns with the biological processes of synaptic development and refinement. Given that synaptic connections initially form abundantly, with many connections later being pruned based on activity and functional relevance, by first learning a dense set of weights with Dale’s backprop, the network can capture a rich set of connections that adhere to Dale’s law, reflecting excitatory and inhibitory roles at a fundamental level. Subsequent application of top-prob pruning mirrors the refinement phase, where weaker, less functionally critical synapses are eliminated, retaining only the most effective pathways. This pruning rule not only emphasizes synaptic efficacy [52] — preserving stronger synapses — but also adheres to principles of synaptic scaling [53] in the retraining phase^2^ by maintaining a balanced level of activity within the network. Overall, this process of initial dense learning followed by selective refinement embodies how biological systems evolve and ensures computational efficiency by optimizing for both anatomical and functional plausibility [54, 55].

We demonstrate the suitability of our methods for studying neuroscientific data by applying them to RNNs trained to fit a two-photon calcium imaging dataset exploring multi-regional interactions that underlie visual behavior in mice when performing a change detection task [56]. We find that our models successfully recapitulate both long- and short-timescale interactions among neuronal populations, capturing transient dynamics as well as sustained signals that are critical for complex perceptual processing. Moreover, our model outputs align with the predictive coding hypothesis [57], as they reflect anticipatory and feedback-driven patterns observed experimentally, suggesting that our approach is well-suited to modeling the layered processing of sensory information in the brain.

Taken together, our results on synthetic and real-world datasets indicate that our methods offer a robust framework for fitting and modeling neural dynamics in a biologically faithful manner. By capturing both anatomically realistic connectivity patterns and functional interactions, our approach provides a set of powerful tools for understanding the complex, hierarchical processing of information across different cell-types, populations, and brain areas. These tools subsequently enable models to better reflect anatomical structures, thereby imparting greater confidence in their findings and enhancing the alignment between RNNs and real neural circuitry.

## 2 Training Networks with Dale’s Backpropagation

In this section, we introduce our sign-constrained learning rule, **Dale’s backpropagation** and validate its performance. In particular, we provide intuition for the algorithm, describe it in detail, present the statements of the theoretical analyses performed (i.e., convergence guarantees and error bounds), and finally provide empirical results demonstrating its utility on a set of neuroscience-inspired and ML tasks.

### 2.1 Dale’s Backpropagation: Algorithm

Dale’s backpropagation enforces Dale’s principle by integrating sign constraints into the conventional backpropagation process. Specifically, it employs a projection step (similar to that of projected gradient descent) on the learnt parameters at every iteration to ensure that the weights remain non-negative for excitatory neurons and non-positive for inhibitory neurons, thus adhering to biological constraints (Fig. 1.B).

Consider the typical Elman RNN [58], whose hidden states *h*_*t*_ at time *t* are updated as per the rule

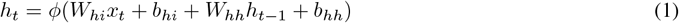

where *ϕ* is the non-linear activation function, *W*_*hh*_ is the recurrent weight matrix, and *W*_*hi*_ is the projection matrix that acts on inputs *x*_*t*_. Biases corresponding to the input and hidden states are denoted as *b*_*hi*_, *b*_*hh*_ respectively.

When *h*_*t*_ are non-negative, respecting Dale’s law simplifies to constraining the recurrent weights *W* such that if *i* is the pre-synaptic neuron and *j* is the post-synaptic neuron,

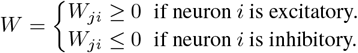

At initialization, the recurrent matrix *W* can satisfy the sign constraints by construction. However, given that standard gradient descent-based backpropagation update

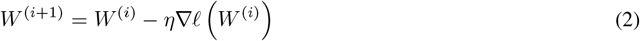

with the step size *η* and loss function *ℓ*, there is no guarantee that the updated weights *W* ^(*i*+1)^ at the next iteration will satisfy the sign constraints set by Dale’s law even if they are respected by the matrix *W* ^(*i*)^ at iteration *i*.

We note however that our sign constraints always form a convex set [59], enabling us to adapt any gradient-based optimization scheme (e.g., SGD, ADAM, RMSprop, etc.) into its projected version [60]. Hence, after the standard backprop update at every iteration, we project the weights onto their feasible set – the orthant in parameter space where the sign constraints of all individual synaptic weights are met. Mathematically, this new update rule can be expressed as

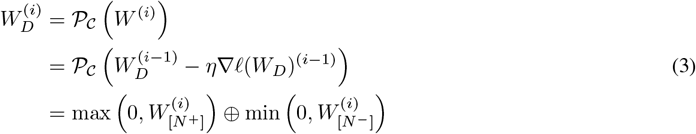

where 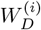 represents the weight matrix that satisfies Dale’s law at iteration *i* and P_*C*_ denotes the projection operator. Weight subsets corresponding to excitatory and inhibitory neurons are denoted by [*N*^+^] and [*N*^−^] respectively. The full derivation of the update is provided in Appendix 6.1. The explicit algorithm for the update under gradient descent as the optimizer is shown in Appendix 6.2.

Moreover, this projection onto the feasible set has both, a simple interpretation and implementation. At every iteration, weights that violate their assigned sign constraints are set to zero, while those that comply are retained at their updated values. The projection itself can be efficiently implemented by multiplying the weights with a binary mask after each update. This flexibility makes it easy to apply our method across various architectures, seamlessly integrating sign constraints within standard backpropagation frameworks.

Consequently, for a single-layer RNN with *N* neurons of which *N* ^+^ are excitatory and *N*^−^ are inhibitory, our entire algorithm for Dale’s backpropagation can be summarized as follows

#### Algorithm 1

Dale’s Backpropagation

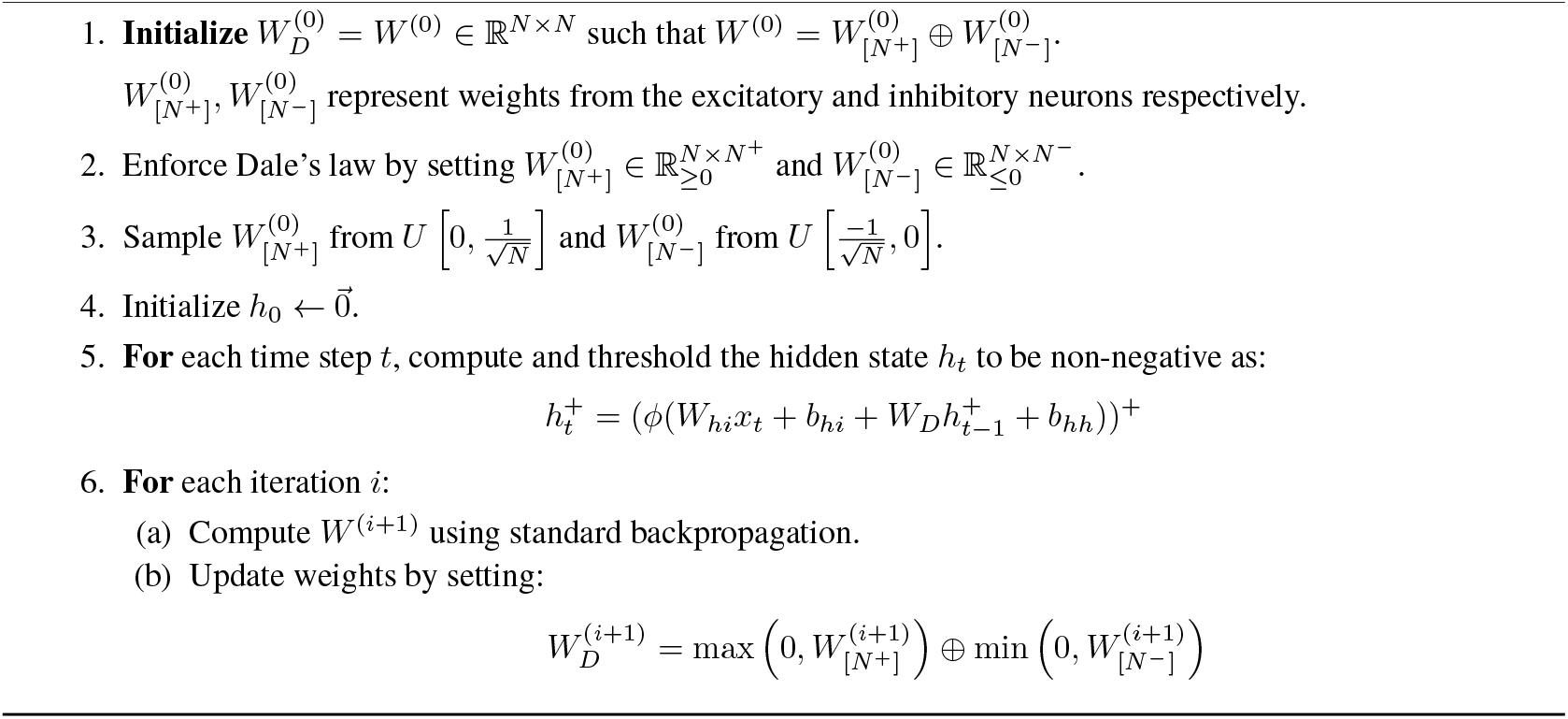

We note that our initialization scheme relates closely to popularly used schemes such as Glorot [61] and He [62] initialization. In particular the scheme aligns with (and is equivalent to when the number of excitatory and inhibitory neurons are equal) common weight initialization practices in RNNs where weights are initialized with zero mean and scaled variance – often using the distribution 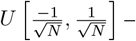 to maintain consistent activation variances, facilitating effective training and convergence [38, 63].

We also observe that thresholding *h*_*t*_ to be non-negative mimics the behavior of biological neurons, in that neuronal firing cannot be negative. Since our goal is to always ensure that *h*_*t*_ is non-negative, we can also increase the threshold to any value greater than 0, depending on the application. Furthermore, it is of interest that previous works were able to enforce sign constraints with FORCE learning [13] in spiking neural networks (SNNs) [47] using an update rule that is the same as ours, but since *h*_*t*_ is non-negative by definition in an SNN, this is not a step that they explicitly needed to incorporate into their training algorithm. Additionally, we can show that despite their different formulations, Dale’s backpropagation reinforces an overlapping set of synaptic connections as Hebbian learning [64], thus preserving key aspects of biologically plausible learning dynamics (Appendix 7).

Finally, it is worth mentioning that the Dale’s backpropagation update is guaranteed to find the weights *W*_*D*_ that are the closest projection of *W* under sign constraints **(Theorem 5)**, with respect to the Frobenius norm (Appendix 6.3). Other methods that are similar to ours ideologically choose to enforce Dale’s law by always using a ReLU activation function and consequently constraining the post-synaptic weights to be positive (or negative) for excitatory (or inhibitory) neurons [50, 45] by multiplying these non-negative weights with a mask comprised of ±1. While this method (which we call *rectified backprop*) is equivalent to ours in the case of weights projecting from excitatory neurons, it essentially reverses the sets of weights that are kept vs. zeroed out in the case of inhibitory neurons. In terms of the final update, the new weights learnt by rectified backprop therefore are much farther away than the one originally computed by conventional backpropagation **(Corollary 6)**.

### 2.2 Dale’s backpropagation: Theoretical Results

We now present our key mathematical analyses for Dale’s backprop when it utilizes gradient descent as its optimizer. First, we derive the rate of convergence of the algorithm under the assumption of restricted optima, i.e., when we can assume that the optimal set of parameters also have the same sign pattern as those imposed. Second, we quantify the differences between Dale’s backprop and standard backpropagation, both in terms of the weights learnt and the final solutions obtained. Together, these results establish a solid theoretical foundation for Dale’s backprop, demonstrating its ability to learn effectively and efficiently, thus validating its use in modeling neural data.

#### 2.2.1 Analyzing convergence of Dale’s backpropagation under the restricted optimum assumption

We start by examining the behavior of Dale’s backpropagation algorithm under the assumption that the optimal set of parameters for a task shares the same sign pattern as the one imposed – a condition we refer to as the *restricted optima assumption* – and show that despite having to learn with constraints, under this assumption, Dale’s backprop converges linearly to the optimal solution **(Theorem 2)**. Biologically, this assumption mirrors the idea that the arrangement of excitatory and inhibitory neurons in the network is optimized for such tasks.

Proving this theorem relies on the geometric observation that the restricted optima assumption ensures that the globally optimal set of weights (*W*^∗^) lies within the same orthant as our point of initialization *W* ^(0)^. We subsequently prove optimal sign pattern preservation **(Lemma 1)**, which guarantees that every backpropagation iteration never leaves this orthant, implying the signs of the weights remain constant throughout the optimization process. As a result, Dale’s backpropagation behaves identically to unconstrained gradient descent within this orthant, making the projection step redundant since the optimization path does not approach the boundaries of the orthant. Consequently, the algorithm can take the most direct path to the optimum without any detours induced by constraint enforcement, allowing it to achieve a linear convergence rate under the Polyak-Łojasiewicz condition.

The statements of our results are presented below, with full proofs deferred to the supplement (Appendix 8.1).

##### Lemma 1

(Optimal sign pattern preservation). *Let the vector of learnt weights be W* ∈ℝ^*n*^ *with the components w*_*j*_, *where j* ∈[1, 2, …, *n*}. *Let L be the Lipschitz constant for the gradients ∇ℓ*(*W*), *where ℓ is a loss function. Given a gradient descent-based, component-wise sign-preserving learning rule that uses the projection operator* 𝒫_*C*_ : ℝ^*n*^ 1→ ℝ^*n*^ *defined as*

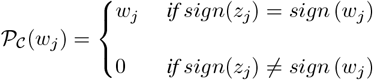

*where* 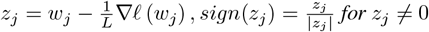, *and sign*(0) = 0. *If sign*(*W*^∗^) = *sign*(*W* ^(0)^) *where W*^∗^ *are the set of weights that can achieve the optimal loss on ℓ, it holds that for any iteration i of regular gradient descent*

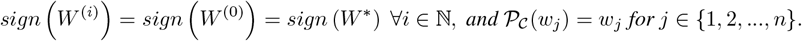

##### Theorem 2

(Convergence of Dale’s Backpropagation). *Let ℓ be a loss function satisfying the µ*−*Polyak-Łojasiewicz condition, with gradients that are L-Lipschitz such that L* ≥*µ >* 0. *Consider the sequence of weights*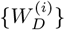 *generated according to the Dale’s backpropagation update, with a step size of* 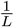. *Given an optimal loss ℓ*^∗^ = *ℓ*(*W*^∗^) = argmin *ℓ*(*W*_*D*_) *where W*^∗^ *has the same sign pattern as all* 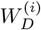 *and a specific error ε >* 0, *it holds for iteration i that*

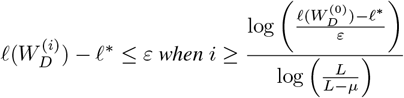

Notably, our analysis reveals that under the assumption of the restricted optima condition, Dale’s backpropagation achieves a linear convergence rate which matches the performance of unconstrained backpropagation [65]. This is significant, as it demonstrates that under the right conditions, imposing biological constraints through Dale’s principle does not necessarily come at the cost of convergence speed. Furthermore, it also suggests that the brain’s neural circuitry despite being constrained by Dale’s law might also be functionally organized to facilitate efficient learning and task performance. However, it is important to note that these guarantees rely not only on the restricted optima assumption, but also on the gradients satisfying Lipschitzness and the Polyak-Łojasiewicz condition, which from a mathematically rigorous perspective may not always hold in practice.

#### 2.2.2 Analyzing Dale’s backpropagation relative to conventional backpropagation

Dale’s backprop also lends itself well to analyzing its behaviour with respect to standard backpropagation when we do not make the restricted optima assumption. Specifically in the case of a single-layer recurrent neural network (without biases), we can characterize the distance between the weights found using standard backprop and Dale’s backprop **(Lemma 3)**, and therefore subsequently the distances between outputs found using the two weight update schema, allowing us to bound the difference between the final error of the solution found using Dale’s backprop in terms of that found using standard backprop **(Theorem 4)**. Formally, we express the above as follows:

##### Lemma 3

(Distance between learnt weights). *Let W* ^(*i*)^ *and* 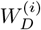 *be the weights at iteration i for standard backpropagation and Dale’s backpropagation, respectively. Assume the gradients* ∇*ℓ*(*W*) *and* ∇*ℓ*(*W*_*D*_) *are upper bounded in magnitude by G and Lipschitz continuous with constant L. Then, the distance between the two sets of weights at any iteration i, denoted as* 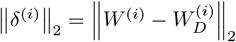 *is bounded*^3^ *by:*

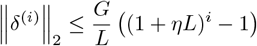

*where η is the learning rate*.

##### Theorem 4

(Differences in errors between solutions). *Let f* (*W*) *be the function represented by a single-layer RNN unrolled over T timesteps, with weights W. Let W*_*D*_ *be the weights learnt using Dale’s backpropagation, and W be the weights learnt using standard backpropagation. Assume the non-linearity ϕ is either tanh or ReLU. Then, the error of the solution found using Dale’s backpropagation with respect to the ground truth y is bounded by:*

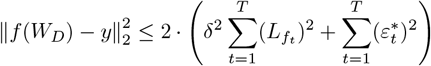

*where f* (*W*_*D*_) *is the output after K training iterations and* 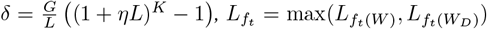 *is the maximum of the Lipschitz constants of the two RNNs at timestep t, and* 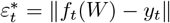 *is the error of the solution found using conventional backpropagation at timestep t*.

The lemma on the distance between learnt weights quantifies how Dale’s backpropagation diverges from standard backpropagation over time due to the sign constraints, showing that this divergence grows but remains bounded, influenced by factors like the learning rate, the loss landscape’s smoothness, and gradient magnitudes. Building on this, the theorem on error between solutions relates the performance of Dale’s backpropagation to standard backpropagation, indicating that the error of Dale’s method is bounded by both the divergence of the weights (bounded by the lemma) and the sensitivity of the network’s output to weight changes, alongside the error of the standard method. Together, these results provide theoretical assurances that, while biological constraints impact learning dynamics, they do not cause uncontrolled error growth, supporting the use of Dale’s principle in neural network training.

Full proofs for both the lemma and theorem are provided in the supplement (Appendix 8.2).

### 2.3 Dale’s backpropagation: Empirical results

We evaluate the performance of Dale’s backpropagation across three tasks of interest (Fig. 2.A). The first is a 1-bit **flip-flop** task (Fig. 2.A - top row - left), in which the network is required to maintain and toggle between different states in response to a series of binary inputs. Specifically, the network output is meant to start at zero, following which it takes the value ±1 to match the input signal whenever presented. It is then expected to switch signs if presented with a signal of the opposing sign, or else, maintain the same output as before. Next, is a **wave reconstruction** task (Fig. 2.A - bottom row) where both the excitatory and inhibitory neurons are presented with individual sinusoidal waveforms. The network is tasked with accurately reconstructing both signals simultaneously, reflecting the roles of excitation and inhibition in modulating distinct aspects of signal processing in neural circuits. We finally also test our methods on the **sequential MNIST** task (Fig. 2.A - top row - right), which is a variation of the classical digit classification task in that instead of receiving the entire image as the input, the network instead sequentially receives the rows of the image.

**Figure 2:**
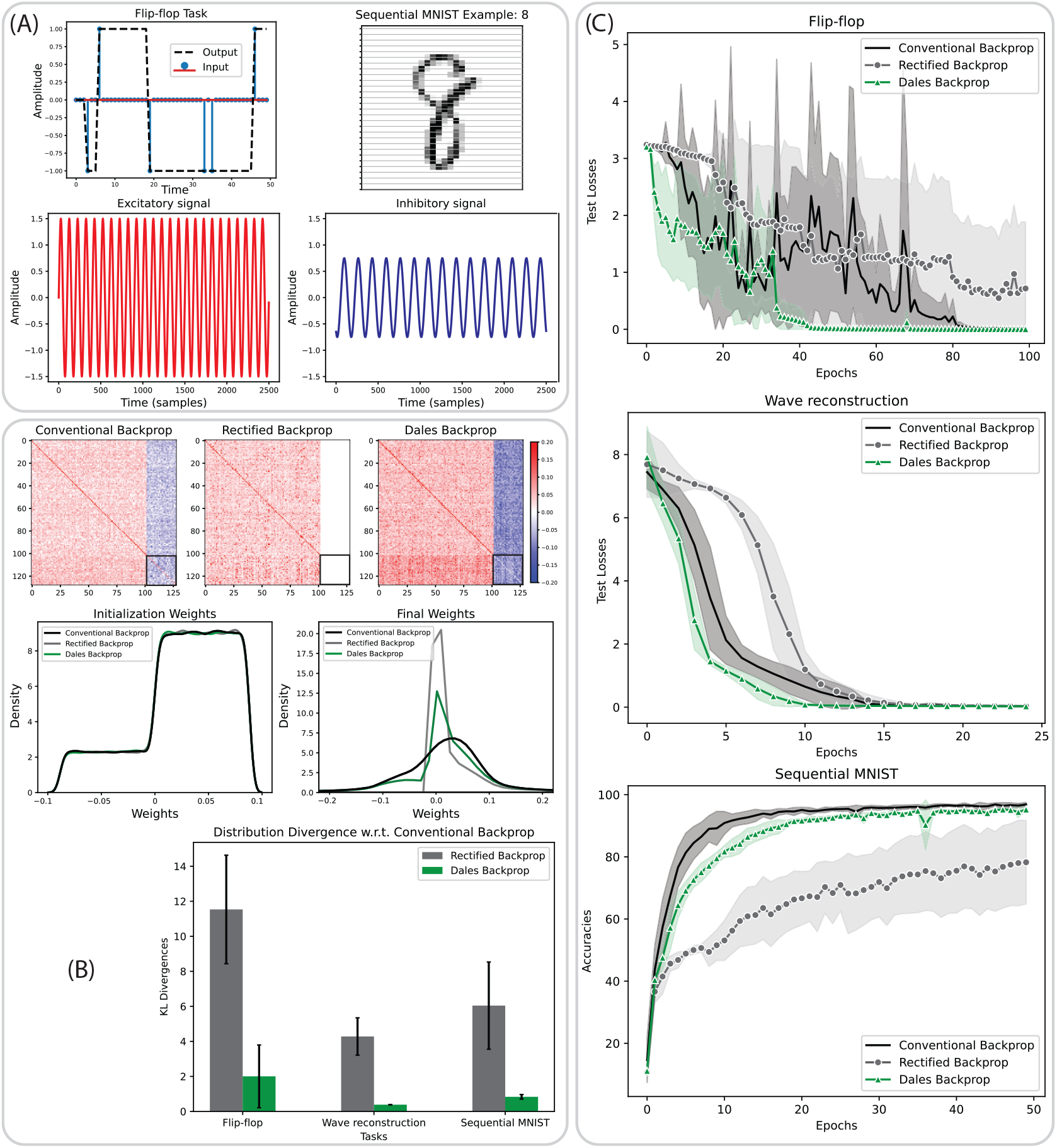
Training with Dale’s Backprop. **(A)** Task examples: 1-bit flip-flop, Sequential MNIST, Wave reconstruction. **(B)** Distribution of weights: Weight matrices post-training (top row), Weight histograms at initialization and after training (middle row), Relative divergence in weight distributions with rectified and Dale’s backpropagation vs. conventional backpropagation (bottom row). **(C)** Test performance of models across different tasks when trained with conventional backpropagation (black), rectified backpropagation (gray), and Dale’s backpropagation (green). All statistics computed over 5 independent runs.

For each of the tasks we train RNNs (over 5 runs) with 128 hidden neurons of which ∼80%(102) are excitatory and ∼20%(26) are inhibitory. In addition to conventional backpropagation, we also include in our experiments “rectified backprop” a similar method from the literature [50, 45] that is also amenable to incorporating sparsity constraints^4^ simultaneously. Our experiments justify our proposed weight update and subsequent theoretical analyses, both in terms of how the weights evolve, as well as learning performance. We start by analyzing the distribution of weights before and after training for the three learning rules (Fig. 2.B) averaged across all tasks. While we initialize all three methods identically (Fig. 2.B - middle row - left), we notice that their distributions post-training are visibly different (Fig. 2.B - middle row - right). Specifically, the weights learnt using conventional backprop (black) show the smallest peak around zero and the greatest deviation, followed by those learnt using Dale’s backprop (green), and finally rectified backprop (gray). This trend is also visible in the weight matrices themselves (Fig. 2.B - top row), where the negative weights learnt using rectified backprop seem practically to be zero (Fig. 2.B - top row - middle). On first glance the weight matrices for conventional and Dale’s backprop seem almost the same, but closer inspection (black boxes – bottom right of the weight matrices) reveals that some of the weights which should have been negative have flipped signs with conventional backprop (Fig. 2.B - top row - left) but there are no such discrepancies with Dale’s backprop (Fig. 2.B - top row - right). Finally, we empirically quantify the differences in learnt weights for the two sign-preserving methods by measuring the Kullback-Liebler (KL) divergence amongst their weight distributions post training (Fig. 2.B - bottom row) with respect to conventional backpropagation. We observe that across all three tasks the divergence shown by rectified backprop (gray) from the weights learnt by conventional backprop are much higher than those shown by Dale’s backprop (green).

Finally, the learning performance (Fig. 2.C) of Dale’s backprop (green) matches that of conventional backprop (black) for all three tasks, at times even learning faster. We conjecture this is a consequence of the regularization introduced by adhering to the sign constraints – by restricting the optimization to the orthant where the signs are preserved, the search space is effectively reduced. This focused parameter space allows for more efficient learning dynamics, as the optimizer concentrates on adjusting weight magnitudes without expending effort on sign changes that violate the constraints. The fixed signs lead to more stable and directed weight updates, resulting in a smoother optimization landscape. Consequently, Dale’s backprop can converge more rapidly in the early stages of training, while ultimately achieving similar final performance as standard backpropagation. However, the learning performance of rectified backprop (gray) is both slower and not as competitive as the other two methods, especially on the more complicated sequential MNIST task. To that end, we note that this might be a consequence of the fact that rectified backprop does not allow for activation functions that have any negative outputs (e.g., tanh). This restriction likely leads to exploding gradients and dead neurons – the latter which might also be inferred visually from its weights post training. However Dale’s backpropagation does not suffer from such limitations and can use non-linearities that have negative outputs as long as it is centered around 0, i.e., it keeps positive and negative activations, positive and negative respectively.

## 3 Sparsifying Networks via Topology-Informed Local Pruning

Having established a method that lets us train sign-constrained RNNs leveraging the machinery of autograd-based backpropagation [66, 67], in this section we describe **topologically-informed probabilistic (top-prob) pruning** as a way of sparsifying dense neural networks to reflect a target connectivity pattern amongst neuronal populations (Fig. 1.C). We first describe the pruning rule formally, while also motivating it from both a mathematical and neuroscientific perspective. Our subsequent empirical analyses demonstrate the applicability of our method in conjunction with Dale’s backpropagation, wherein it outperforms the random pruning baseline on different tasks.

### 3.1 Topologically-informed probabilistic pruning rule

Consider a weight matrix *W* ∈ ℝ^*N×N*^ comprised of the synaptic weights *w*_*ji*_ ∀*i, j* ∈ [1, 2, 3, …, *N*}, connecting neuron *i* → *j*. The sparsified matrix *W*^*sparse*^ ∈ ℝ^*N×N*^ is obtained using the pruning rule

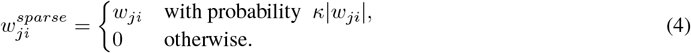

where *κ* ∈ ℝ^+^ is a non-negative scalar that controls the sparsity of the resulting matrix and is defined as

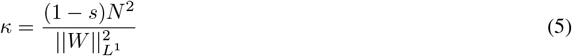

*s* ∈ [0, 1] is the target sparsity of *W*^*sparse*^ and 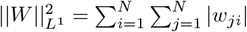.

While it is evident that the top-prob pruning rule operates by probabilistically retaining weights of higher magnitude while eliminating weaker ones, we emphasize that this approach mirrors fundamental aspects of synaptic plasticity in biological neural networks. The rule’s local nature – where pruning decisions depend solely on individual synaptic weights – aligns with biological constraints, as real neurons modify their connections based only on local synaptic properties rather than global network states. Furthermore, when coupled with Dale’s backpropagation, the top- prob pruning mechanism has a propensity for preserving exactly those weights that align with Hebbian learning principles (Appendix 7), thereby ensuring the maintenance of biologically meaningful functional connectivity whilst simultaneously achieving network sparsification.

The top-prob approach is also grounded from an ML and mathematical standpoint. Magnitude-based pruning has a long history [48, 68, 69] and in its iterative form is still a highly competitve empirical baseline for neural network compression via pruning [39, 40]. It also closely relates to methods that look to preserve weights that maintain the dynamics of the network in the spectral sense [49, 70, 71, 72]. Finally, it provides us with an elegant way of maintaining the structural integrity of the network. Recent works have established that the zeroth-order topological information of a graph is fully encapsulated by its maximum spanning tree (MST)^5^ [74, 75, 76, 73]. Ergo, probabilistically maintaining higher magnitude weights of the network results in us preserving this topological structure (Appendix 10.1).

Throughout the remainder of this work, we use top-prob pruning in the one-shot sense [41], wherein we prune to a target sparsity level and connectivity pattern in a single step, followed by a single retraining phase to help restore the model’s performance. We note however that our approach can just as easily be used at initialization to sub-select a sparse network pre-training, or alternatively used in the iterative manner that is more typical in the ML community, especially for tasks that are more complicated and less amenable to drastic drops in sparsity levels from a fully-trained dense configuration. Additional explanations for how we derive and re-normalize our hyper-parameter *κ* across different contingencies, as well as adjust the parameter *s* are provided in Appendix 9.1, 9.2, and 9.3 respectively.

### 3.2 Topologically-informed probabilistic pruning: Empirical results

We study the behaviour and performance of top-prob pruning by first examining how it impacts the weight distribution and structural integrity of the original dense network, followed by its ability to maintain functional capacity. Our results suggest that top-prob pruning does indeed preserve key network properties, leading to highly sparse yet robust models that do not require extensive retraining to regain performance.

As a preliminary check we observe the distribution of non-zero weights (Fig. 3.A) for dense weight matrix (black), vs. that of a matrix that has been pruned to 90% sparsity using the top-prob pruning rule (green) and random pruning (grey). As expected, we see the dense matrix has weights that are almost uniformly distributed since the weights were sampled from the distribution 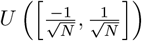 and the randomly pruned network stays faithful to this distribution. The weights retained using top-prob pruning however are heavily skewed towards being higher in magnitude, demonstrating that it successfully prioritizes stronger connections while eliminating weaker ones. We subsequently quantify the notion of “structural integrity preservation” by plotting the fraction overlap of retained weights with the MST of the dense matrix as sparsity increases (Fig. 3.B) for both pruning methods. Again, we observe that empirically top-prob pruning shows a much higher overlap with the MST (green dots) compared to random sparsification in practice (grey crosses) and theoretically (black crosses) (Appendix 10.2). Additionally, it shows lesser variations in the amount of overlap as well.

**Figure 3:**
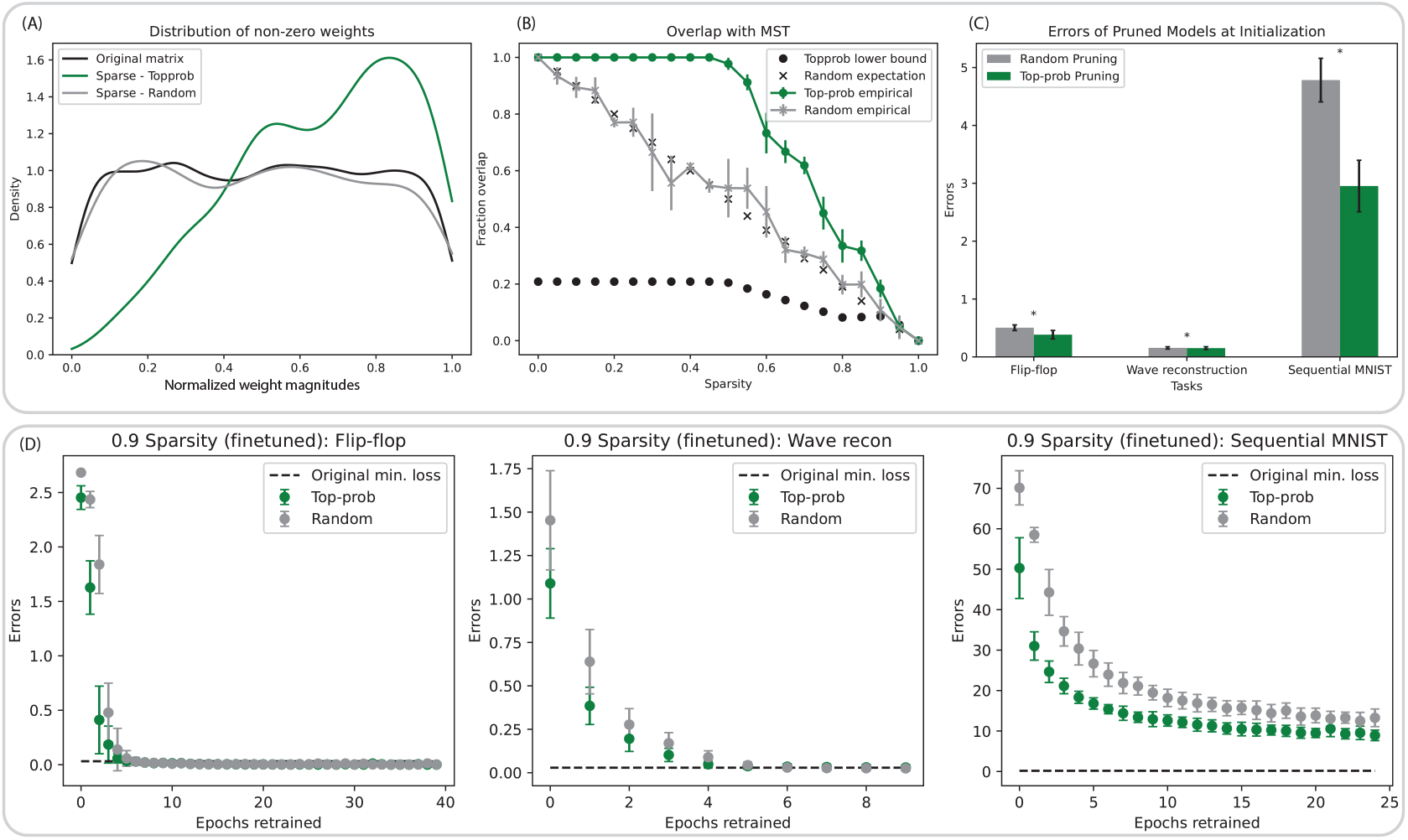
Topologically-informed probabilistic pruning. **(A)** Distribution of non-zero weights for dense (black), and sparse matrices pruned randomly (grey) and with top-prob pruning (green). **(B)** Fraction overlap that retained weights have with the MST of the dense matrix. **(C)** Errors of pruned models, without any retraining using random (grey) and top-prob pruning (green). **(D)** Performance of sparsified and fine-tuned models across different tasks, when pruned randomly (grey) vs. top-prob pruning (green). All statistics computed over 5 independent runs. ^*^ indicates *p* ≤ 0.05.

Moving on to the pruned network’s ability to retain information and functional capacity, we note that the errors post-pruning but before fine-tuning (Fig. 3.C) are always higher for models that are pruned randomly (grey) vs. those pruned with the top-prob pruning rule (when pruned to 90% sparsity). Moreover, the difference in errors becomes more significant as the task complexity increases (p = 0.022 for the flip-flop and wave reconstruction tasks, while p = 0.012 for sequential MNIST). Fine-tuning with Dale’s backpropagation for ∼50% the number of epochs as the original training, while retaining the sparse structure identified via pruning follows a similar trend (Fig. 3.D), with networks that that were pruned randomly (grey) showing higher errors at the epoch of re-training than those pruned with top-prob pruning (green). While both methods lead to models that seem to eventually approach the original model’s performance (dashed line) after fine-tuning, top-prob pruning consistently starts from a better initial error, converges faster to optimal performance, and shows more stable learning. We therefore conclude that our pruning rule effectively identifies and retains functionally important weights. The preserved connectivity aligns well with the network’s core topology (MST) which leads to efficient and robust performance of the sparsely structured RNN, in conjunction with Dale’s backpropagation.

## 4 Application to Visual Behaviour in Mice: Functional Connectivity and Predictive Coding

Having established the efficacy of our methods in successfully constructing and training highly sparse RNNs which respect Dale’s law, we apply them to study visual behaviour in mice under the predictive coding hypothesis [57, 77]. Specifically, we model data from the Allen Institute Visual Behavior dataset [78, 79], which comprises two-photon (2p) calcium imaging recordings from mice performing a change detection task, when presented with expected and unexpected stimuli. This experimental paradigm allows us to investigate how different neuronal populations in the cortical circuit interact and process information under varying predictive contexts, shedding light on how prediction errors may be communicated across hierarchically-related cortical regions. Moreover, since our modeling framework captures these interactions while respecting anatomical connectivity and signaling constraints, our analyses reveal how functional connectivity between populations adapts to the inherent biological scaffolding to support such predictive processing, providing insights into the circuit-level implementations of predictive coding in the visual cortex.

In the following subsections we provide details of the specific experimental setup and curated dataset, followed by our model architecture and training methodology. Our results align well with previous observations made in the experimental literature studying the data, and strongly support the predictive coding hypothesis. Furthermore, by “learning” the functional connectivity under various conditions, our approach not only corroborates previous experimental results, but also gives us a way to generate new hypotheses about how different prediction violations engage distinct patterns of feedforward and feedback connectivity across cortical layers and cell types, offering novel insights into the principles governing cortical circuit organization in predictive processing.

### 4.1 Dataset and experimental setup

The Visual Behavior Dataset [78, 79] entails a visually-guided, go/no-go task where mice are shown a continuous series of briefly presented natural images and they earn water rewards by correctly reporting when the identity of the image changes [80]. Responses from the mice are collected as they are presented with two different sets of images; A **familiar set** (Fig. 4.B - top row) comprising images that they were trained on, and a **novel set** (Fig. 4.B - bottom row) that are only presented at test time, during the recordings. While the trials themselves are longitudinal spanning multiple image changes, we restrict ourselves to modeling two full image presentations (Fig. 4.C - Top). If the identity of the second image is the same as that of the first one, we refer to the condition as **no change** (Fig. 4.C - second row). If the identity of the second image is different from that of the first one, we refer to it as the **change** condition (Fig. 4.C - third row). Both images are always from the same set, i.e., they are both either familiar or novel. In a small subset of the trials (∼5%), the second image is omitted, and instead replaced by a blank screen (Fig. 4.C - first row) allowing for analysis of expectation signals. We call this the **omission** condition. For more details on the experimental setup, see [79].

**Figure 4:**
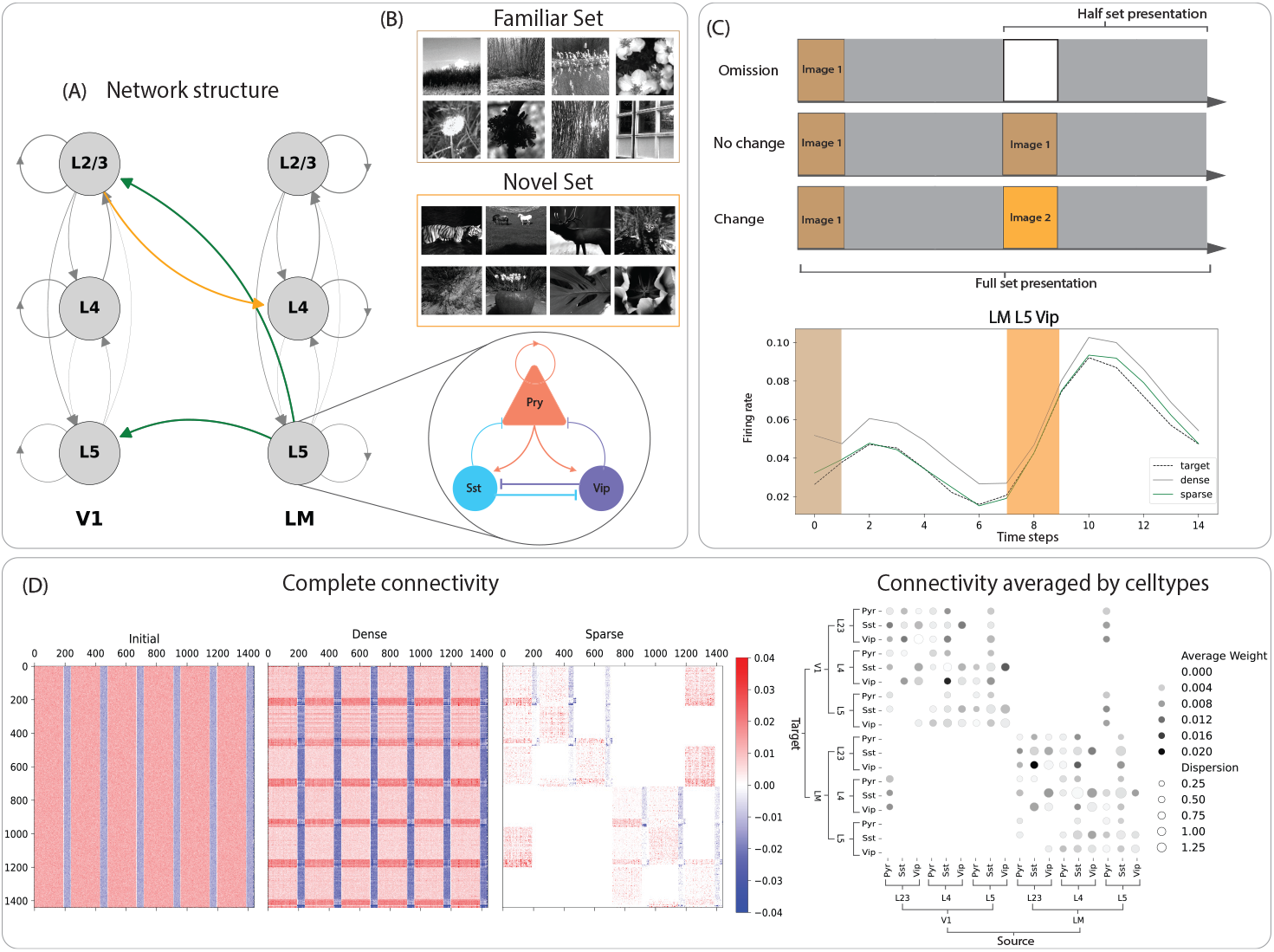
Dataset, network structure, and task schematics. **(A)** General architecture of the CelltypeRNN. **(B)** Familiar and Novel image sets used for training mice on the visual change detection task. Reproduced from the Allen Institute Visual Behavior-2p dataset (open source) [78]. **(C)** Top - Examples of different stimuli conditions in the visual change detection task, depiction of full and half set presentation timescales. Bottom - An example of target activity (dashed black curve), dense RNN output (solid grey curve), and sparse RNN output (solid green curve) from LM L5 Vip population. **(D)** Examples of inferred functional connectivity: Left - Complete neuron-to-neuron connectivity at initialization, after training with Dale’s backprop, and after sparsification (and retraining) with top-prob pruning. Neurons are ordered by area (V1 followed by LM) within which they are ordered by layer (L4, L2/3, L5), and type (Pyr, Sst, Vip). Right - Example of the sparse connectivity matrix where activity is averaged by cell type in every layer (bigger circles imply higher dispersion and darker colours imply stronger connections; dispersion is computed as the fraction of standard deviations to the mean activity in the population).

For each of our conditions we consider two temporal windows. In the **full-set presentation** (Fig. 4.C - indicated at the bottom), we model neural activity across the entire two-image sequence (first image (250ms), inter-stimulus interval (500ms), second presentation/omission (250ms), and post-stimulus interval (500ms)), which allows us to capture the <u>sustained dynamics underlying predictive computation across time</u>. In contrast, the **half-set presentation** (Fig. 4.C - indicated at the top) models neural activity following the second presentation/omission, enabling us to isolate the transient neural responses that implement the mechanistic components of prediction and error signalling. This complementary approach provides insights into both, the overarching dynamics and the immediate neural interactions that support predictive coding, and gives us the flexibility to infer both long-term and short-term functional interactions.

The complete dataset includes multi-regional 2-photon data from *two hierarchically adjacent areas*, **VISp** (i.e., primary visual cortex or **V1**) and **VISl** (i.e., the lateromedial area or **LM**). For both areas we collect recordings at *depths* roughly corresponding to **layers 2/3, 4, and 5** in the cortical column for *excitatory*, i.e., pyramidal **(Pyr)** neurons and *two types of inhibitory neurons*, viz. somatostatin **(Sst)** and vasoactive intestinal peptide **(Vip)** expressing interneurons (sampling depths distributions provided in Appendix 11). In total, we therefore model the activities of <u>18 different interacting populations</u> (Fig 4.A).

To curate the training data for our RNNs, we compute the neuron-averaged response for every experiment corresponding to each of our individual neuronal populations (e.g., LM L5 Vip) from the Allen Institute Visual Behavior-2P dataset [78]. We then randomly sample (with replacement) 100 averaged responses from the total set of averaged responses, take their mean, and pass the same through a 1-D Gaussian filter (*σ* = 1) to produce a single training sample (Fig. 4.C - dashed black curve). We subsequently produce 2000 such samples for each of our individual neuronal populations.

### 4.2 CelltypeRNN: Architecture and training

We model the data as described previously with the anatomically constrained **CelltypeRNN** that replicates the inter-areal structure of the canonical cortical microcircuit with two hierarchically related cortical areas (Fig. 4.A) [81, 82, 83, 84] whilst simultaneously enforcing intra-areal lateral connectivity among different cell types within the cortical column as established by [85]. Moreover, given that the CelltypeRNN is constructed to be able to replicate experimentally obtained response patterns in different cell populations as specified by their cell type, cortical layer and area, by learning the connection weights, we in turn represent the inferred functional interactions between the populations across the cortical circuit [20, 11] under different stimulus conditions.

Subsequently, we first train a dense, unbiased Elman RNN using Dale’s backpropagation, following which we prune the network’s recurrent connections block-wise with top-prob pruning (Fig. 4.D, left) to achieve their individual target connectivity sparsities. We subsequently fine-tune the post-pruning non-zero RNN weights to achieve an overall performance that is at least as good as that of the RNN pre-pruning (Fig. 4.C, bottom). In our specific instantiation, the ratio of Pyr:Sst:Vip neurons in every layer is 12:2:1 (making the excitatory:inhibitory neuronal ratio 4:1), which with a scaling factor of 16 gives us a total of 240 neurons per layer and 1440 overall in the model. Our lateral connectivity probabilities across populations follow experimental data [85] and are explicitly stated in Appendix 12. Longer range inter-areal projections are sparsified to have a connection probability of 0.3, and are strictly excitatory, i.e., Feedforward connections: V1 L2/3 Pyr → LM L4 Pyr, Sst, Vip. Feedback connections: V1 L2/3 Pyr, Sst, Vip ← LM L5 Pyr and V1 L5 Pyr, Sst, Vip ← LM L5 Pyr.

In addition to the weights of the RNN – i.e., input weights *W*_*hi*_ and recurrent weights *W*_*hh*_ – we also have readout weights that project the recurrent RNN activity of individual neuronal populations onto their respective output space, using randomly initialized, fully connected linear layers. The readout weights are frozen at the time of initialization of the dense RNN itself, and remain so throughout the training procedure. By doing so we ensure that any changes in the model’s behavior come from changes in the recurrent dynamics, and not the model “cheating” by simply adjusting its output mapping. It subsequently also makes it easier to interpret and compare how the internal representations and computations change across conditions. To that end, we also mask the input weight matrix *W*_*hi*_ so that recurrent neurons corresponding to a specific population do not receive inputs from any other populations.

Our training objective requires each individual population to be able to reconstruct its activity predictively one timestep into the future (Fig. 4.C, bottom), giving us the loss function

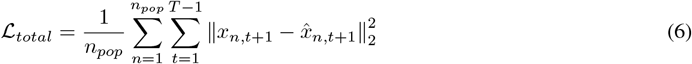

where *n*_*pop*_ is the number of interacting neuronal populations and *T* is the total number of timesteps in the sequence. *x*_*n*,*t*+1_ is the input that will be received for population *n* at timestep *t* + 1 while 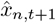 is that predicted by the RNN. The loss function is kept the same during both, the dense training (Fig. 4.C, bottom - grey curve) and fine-tuning post-pruning stages (Fig. 4.C, bottom - green curve). However, we fine-tune for only half the number of epochs (50) as we train for with the dense network (100).

We train separate models for each of our twelve different conditions (Familiar/Novel × Change/No Change/Omission × Full Set/Half Set presentation) and compare their connection weights across various spatial scales, the results of which are discussed in the following subsection.

Our codebase to download and pre-process the data, as well as construct and train the celltypeRNN models across various conditions and timescales is made publicly available at https://hchoilab.github.io/biologicalRNNs.

### 4.3 Insights and results

Using the anatomically-constrained CelltypeRNN architecture, we examine how distinct cell types across the layers and hierarchy in the visual cortex communicate both expected and unexpected information by comparing inferred connectivity patterns across different experimental conditions and timescales. By fitting neuronal responses of interacting populations through one-step-ahead predictive modeling, we capture the dynamic temporal dependencies inherent in neural activity and the RNN’s resulting connectivity matrix serves as a functional proxy for interactions amongst populations, reflecting how signals propagate within the cortical network. Analyzing how the RNN adjusts its connectivity across varying predictive contexts and timescales provides insights into circuit-level implementations of predictive coding, particularly in prediction error communication and modulation of feedforward and feedback pathways over both, entire stimulus sequences and immediate neural responses to prediction confirmations or violations. Our results can be broken down into three key comparisons:

#### Familiar No Change vs. Familiar Change (Full-set Presentation)

In the full-set presentation of familiar images, we observe significant differences in the inter-areal feedforward and feedback connections (Fig. 5.A). When the activities are averaged across layers, there is a stark increase in the projection V1 L2/3 → LM L4 when there is a change in the image compared to when there is not, suggesting that the expectation violation causes enhanced forward communication from V1 to LM (Fig. 5.A - left, middle). Likewise, feedback projections V1 L2/3 ← LM L5 and V1 L5 ← LM L5 are strengthened as well in the change case (Fig. 5.A - left, middle). Even at the scale of cell-types, we observe that the change condition leads to an increase in functional connectivity for both inter-areal feedforward and feedback projections (Fig. 5.A - right, magenta boxes). Additionally, we note that the changes are predominantly red, i.e., the inferred weights in the change condition are generally higher than that in the case of expected stimuli and conditions being perceived (5.A - middle). This trend also holds when we compare familiar and novel stimuli (Appendix 13, Fig. 7) in both the change and no change cases, i.e., the introduction of novelty leads to increased inter/intra area connectivity. In agreement with previous literature [86], this suggests that novelty and unexpectedness increase the brain’s excitability, which in turn could facilitate plasticity and aid learning.

**Figure 5:**
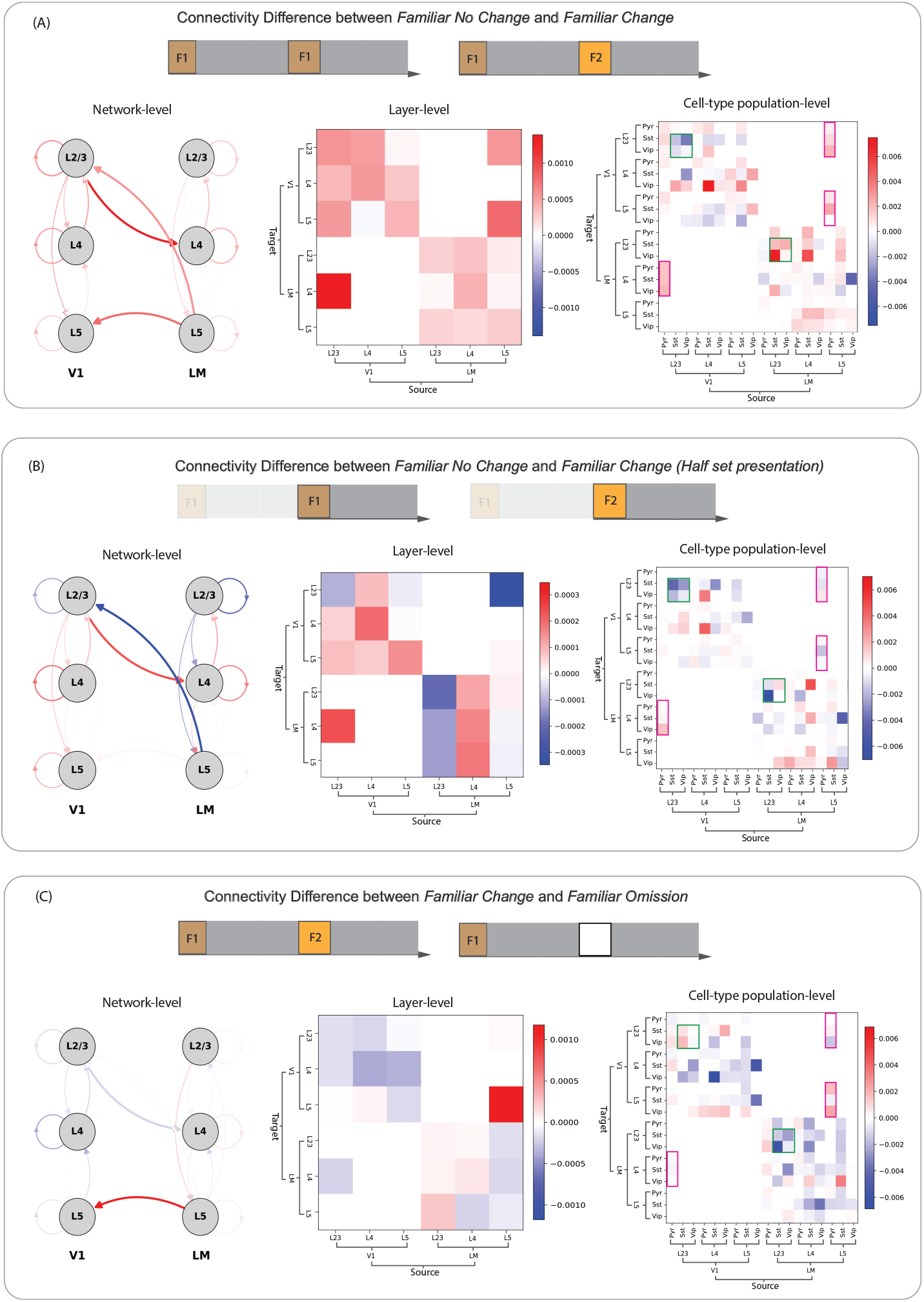
Connectivity differences across timescales and test conditions. **(A)** Familiar No Change vs. Familiar Change (Full set presentation). **(B)** Familiar No Change vs. Familiar Change (Half set presentation). **(C)** Familiar Change vs. Familiar Omission (Full set presentation). All differences are computed as Second condition - First condition; Blue implies higher weights in the first condition, while red indicates higher weights in the second. In all three cases, the left and middle plots are a graphical representation of the weights averaged across layers, while the rightmost plot averages weights by cell-type within each layer. Magenta boxes highlight the feedforward and feedback connections, i.e., those originating at V1 L2/3 and LM L5 respectively. Green boxes highlight all Sst-ViP interactions in L2/3 of both V1 and LM.

**Figure 6:**
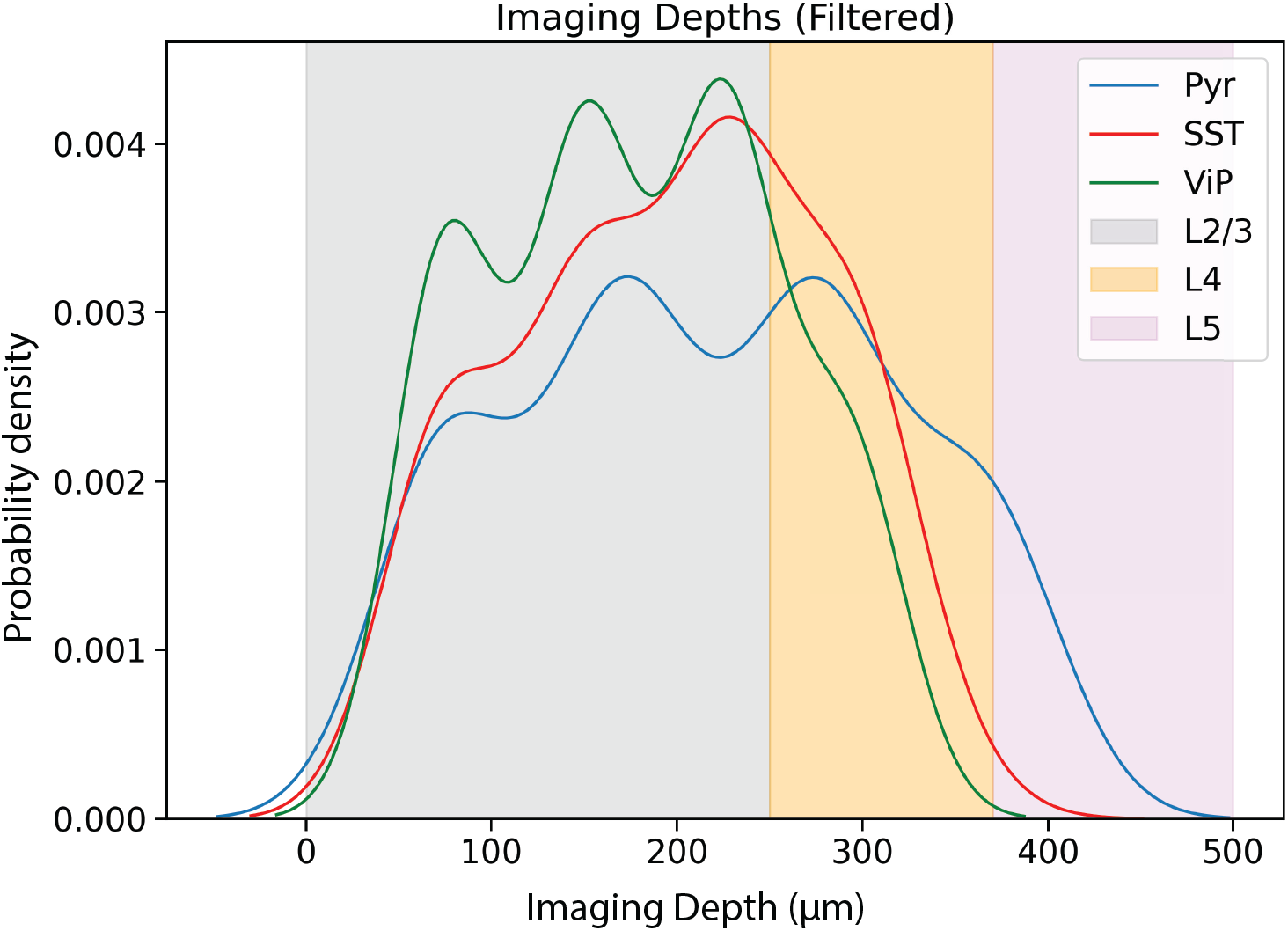
Imaging depths. Distribution of experiments (Y-axis) with respect to imaging depth (X-axis). Shaded grey, yellow and pink panels represent depths corresponding to cortical layers 2/3, 4, and 5 respectively.

**Figure 7:**
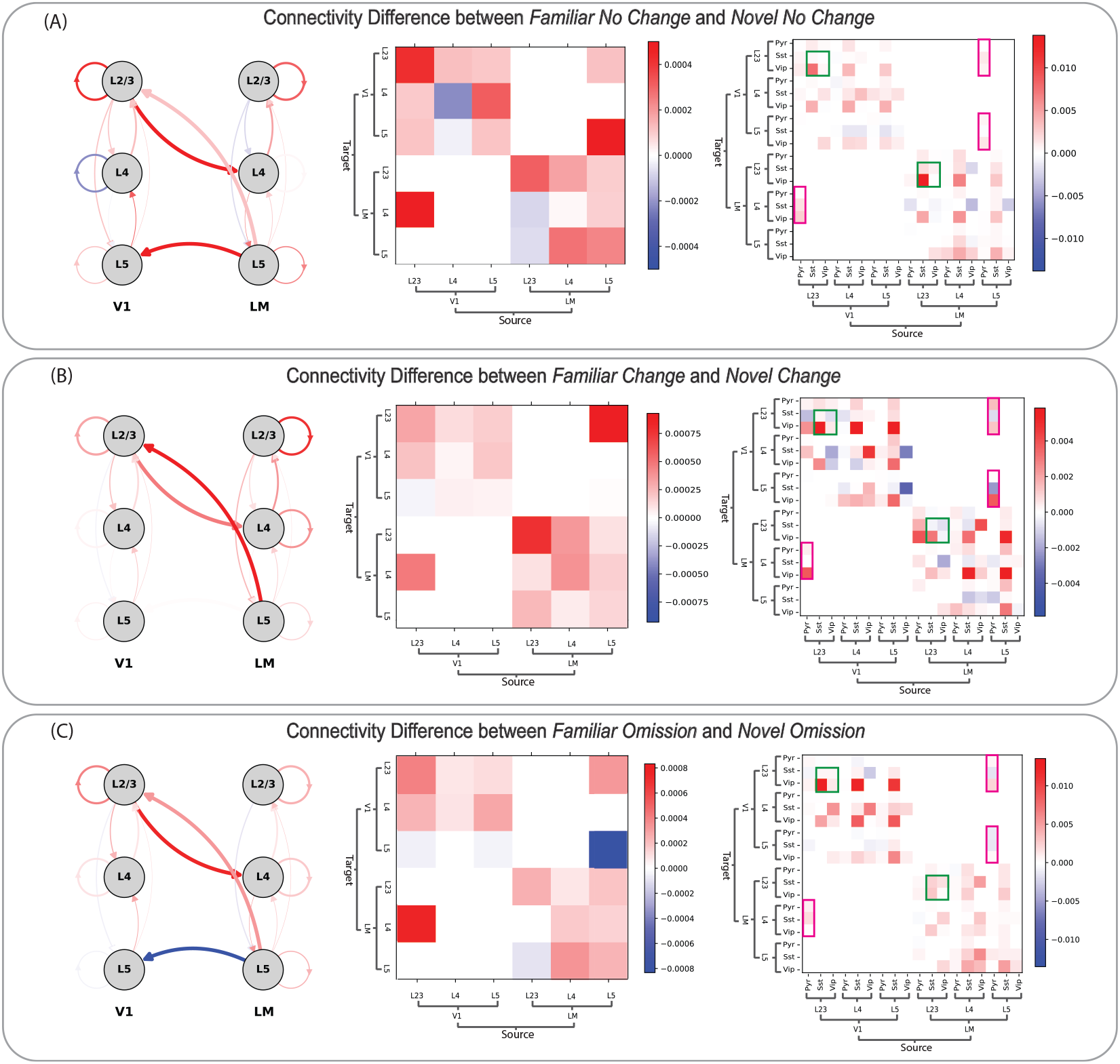
Connectivity differences between familiar and novel conditions. **(A)** Familiar no change vs. Novel no change. **(B)** Familiar change vs. Novel change. **(C)** Familiar omission vs. Novel omission. All plots are from the full presentations condition. All differences are computed as Second condition - First condition; Blue implies higher weights in the first condition, while red indicates higher weights in the second. In all cases, the left and middle plots are a graphical representation of the weights averaged across layers, while the rightmost plot averages weights by cell-type within each layer. Magenta boxes highlight the feedforward and feedback connections, i.e., those originating at V1 L2/3 and LM L5 respectively. Green boxes highlight Sst-ViP interactions in L2/3 of V1 and LM.

#### Familiar No Change vs. Familiar Change (Half-set Presentation)

When focusing on the half-set presentation for familiar images however, we found a contrasting pattern of connectivity with the feedback signaling (Fig. 5.B - left, middle). While we see almost no difference in the weights V1 L5 ← LM L5 across the change and no change cases, the weights V1 L2/3 ← LM L5 are distinctly higher in the *no change* case than the change case. The feedforward projection V1 L2/3 → LM L4 however still maintains the same trend as the full presentation case, wherein it is higher when an image change occurs than when it does not. This suggests that while the feedforward communication is relatively immediate using shorter timescales, propagating feedback information occurs over a longer timescale [83, 87, 88], making a case for further investigation of role of inter-areal, cortico-cortical time-delays [89] when studying predictive coding [84]. The cell-type-specific analysis further reveals that Vip neurons in L2/3 are less inhibited by Sst neurons of the same layer in the change case in both V1 and LM (Fig. 5. B- right, green boxes), compared to the full presentation case (Fig. 5. A - right, green boxes), once again speaking to the transience of the change and also supporting the idea that Vip neurons could encode unexpectedness as hypothesized by the predictive coding theory.

#### Familiar Change vs. Familiar Omission

Our setup also allows us to compare how the network processes different types of expectation violations, by contrasting the learnt weights in the case of an image change vs. image omission. In the setting of familiar images over the length of a full set presentation (Fig. 5.C), we notice that while most of the weights are quite similar for both types of expectation violations, there is an appreciable increase in the feedback projection V1 L5 ← LM L5 in the case of image omission (Fig. 5.C - left, middle). Additionally, this increase seems to be driven by an increase in the feedback connection weight to the Vip cells in V1 L5^6^ (Fig. 5.C - right). These observations are in agreement with experimental findings that omissions trigger signaling in Vip neurons in V1 [90]. We also note that across the RNN, weights to the Vip neurons from locally adjacent Sst neurons are reduced in the omission case, suggesting that these neurons are not as inhibited during the processing of omissions, thus potentially emphasizing their role in processing prediction violations. In the supplement, we also provide results studying the No Change vs. Omission case with both familiar and novel images (Appendix 13, Fig. 8) for fair comparison.

**Figure 8:**
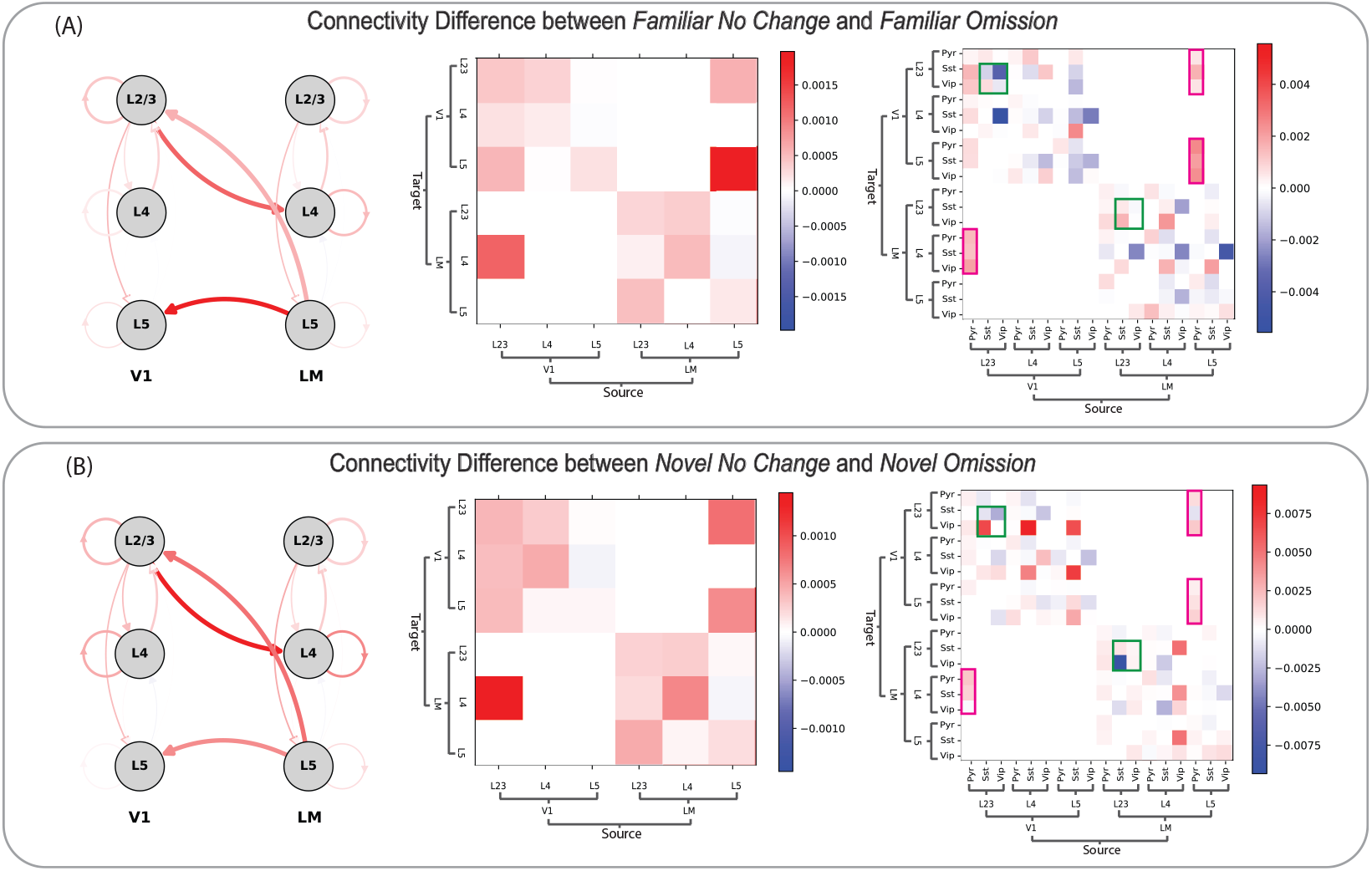
Connectivity differences between no change and omission conditions. **(A)** Familiar No Change vs. Familiar Omission. **(B)** Novel No Change vs. Novel Omission. Both plots are from the full presentations condition. All plots are from the full presentations condition. Differences are computed as Second condition - First condition; Blue implies higher weights in the first condition, while red indicates higher weights in the second. In both cases, the left and middle plots are a graphical representation of the weights averaged across layers, while the rightmost plot averages weights by cell-type within each layer. Magenta boxes highlight the feedforward and feedback connections, i.e., those originating at V1 L2/3 and LM L5 respectively. Green boxes highlight Sst-ViP interactions in L2/3 of V1 and LM.

Collectively, our analysis demonstrates a hierarchical organization of predictive processing in the visual cortex operating over different timescales. We find that feedforward projections are consistently enhanced during all prediction violations across both long and short timescales, emphasizing their crucial role in transmitting prediction error signals (and fundamentally driving synaptic plasticity in the brain [91, 92, 93], facilitating learning and adaptation). In contrast, feedback projections are modulated by both the type of prediction error and the temporal window over which neuronal responses are modeled. Notably, the behavior of feedback projections differs when targeting different cortical layers: feedback projections to L2/3 are more prominently modulated during unviolated predictions over shorter timescales (Fig. 5.B), while feedback projections to L5 are more responsive during negative prediction errors such as omissions of expected visual input [94] (Fig. 5.C). This differential modulation suggests that while feedforward pathways rapidly convey unexpected sensory information, feedback pathways adjust more selectively based on the context, timing, and targeted cortical layer of the prediction error. These patterns are further corroborated by our observations comparing change, no change, and omission across familiar and novel conditions during the full-set presentation (Appendix 13, Fig. 7). In particular, we note that the presentation of a novel image always increases the feedforward connectivity (and ergo the projection) from V1 L2/3 → LM L4. On the finer-scale level of cell-types instead of entire layers, there is consistent prominent involvement of Vip interneurons during prediction violations^7^ which highlights their critical role in modulating cortical circuits in response to unexpected stimuli. Overall, our findings therefore provide circuit-level evidence supporting the predictive coding framework, illustrating how the brain dynamically adjusts its functional neural connectivity in response to varying predictive contexts, timescales, and cortical layers. The dynamic interplay of feedforward and feedback mechanisms facilitates efficient processing of sensory information, enabling the brain to anticipate and adapt to constantly changing environments.

Results for all comparisons across both the full set and half set presentations are publicly available at the project website.

## 5 Discussion

Our work develops methods for constructing RNNs that simultaneously incorporate two fundamental biological constraints: *Dale’s law* and *structured sparse connectivity motifs*. We provide mathematical grounding for these methods, including convergence guarantees and error bounds, demonstrating that they can match the performance of unconstrained RNNs. Empirical results on standard synthetic tasks support the efficacy of our approach, demonstrating that our biologically constrained RNNs can achieve performance comparable to conventional, unconstrained networks. Furthermore, by aligning computational models more closely with biological reality, we enhance their utility for neuroscientific research, providing tools for more accurate modeling of neural dynamics and brain function.

Our approach also differs significantly from CURBD [20], an existing method in the literature for inferring multi-regional interactions, in two key aspects. First, while CURBD successfully models neural dynamics and iteractions, it does not incorporate sign constraints during training, limiting its ability to differentiate between excitatory and inhibitory cellular mechanisms. Second, and more critically, CURBD’s reliance on FORCE training makes it poorly suited for implementing experimentally-informed sparse connectivity patterns among neuronal populations. Every iteration with FORCE is a least-squares update that is dense and doesn’t respect the sparsity constraints of the matrix at the previous iteration - it is non-trivial to subsequently enforce the sparsity pattern, or alternatively solve a recursive least-squares update for every sub-matrix defined by the sparsity pattern at each update, which quickly becomes computationally infeasible. These limitations consequently motivated our development of a our backpropagation-based weight update method that efficiently handles both Dale’s law constraints and structured sparsity.

Applying our methods to the Allen Institute Visual Behavior dataset, we inferred multi-regional neuronal interactions underlying visual behavior in mice performing a change detection task. Our anatomically and physiologically constrained *celltypeRNNs* not only replicated the experimental data but also provided insights consistent with the theory of predictive coding. Specifically, the models revealed dynamic interplay between feedforward and feedback mechanisms across cortical layers and cell types, capturing how the brain adjusts functional neural connectivity in response to varying predictive contexts and timescales.

We note that much of our methodological work can easily be extended to other deep architectures, and is not in fact restricted to simply RNNs. That said, a key area for incorporation of additional biological realism would be in the way we inherently solve the credit assignment problem. Backpropagation suffers from needing a global error signal and weight symmetry [95], prompting the need for more biologically plausible learning rules that can still learn as effectively. One hypothesis is that using local learning rules may contribute to the emergence of more modular network representations by promoting the formation of localized activity clusters, thus leading to deeper insights into how functional specialization arises in neural systems and its role in facilitating learning.

Furthermore, our findings highlight differential neural responses to different types of prediction errors, emphasizing the importance of the nature of the violations in shaping neural dynamics. In our study, the change in the familiar image case represents a “global oddball” – an unexpected stimulus that violates established patterns while maintaining the local context. Conversely, the omission of an expected stimulus constitutes a “local oddball”, introducing a novel scenario for the network. This distinction is significant, as recent work [96] has found that global oddballs elicit responses in non-granular layers, differing from local oddballs that evoke early responses in superficial layers 2/3, consistent with conventional predictive coding theory. Our findings align with this pattern for global oddballs but present discrepancies in the case of local oddballs (omissions). This underscores the need for further exploration into stimulus dependency in error encoding [97], and suggests that normative predictive coding computations may need to account for the type of prediction error to fully capture neural processing dynamics.

Finally, we note that an important consideration in our study is the limited scope of recorded celltypes and brain regions, which poses challenges in interpreting our results. Specifically, we do not have recordings from all interacting celltypes and areas that may be involved in the visual processing tasks we modeled. This limitation means that our models might capture neural responses that are more correlational rather than causal, as they are based solely on the observed data from recorded populations. The absence of data could lead to incomplete or biased representations of neural interactions, especially at the finer grained level of celltypes as opposed to the coarser level of layers, where the absence of a particular subpopulation’s influence is more easily subsumed within aggregate dynamics. To address this gap, future work could involve developing methods that account for unobserved interactions, perhaps through incorporating prior knowledge of anatomical and functional connectivity or using computational techniques to infer missing information. Additionally, expanding experimental recordings to include more brain areas and cell-types would provide a more comprehensive dataset, enabling our models to capture the full complexity of neural dynamics and leading to more causally robust conclusions.

## Acknowledgments

We thank Anqi Wu for insightful comments and feedback. This work was supported by the Alfred P. Sloan Foundation Fellowships in Neuroscience (to H.C.) and the National Eye Institute of the National Institutes of Health under Award Number R00 EY030840 (to H.C.). The content is solely the responsibility of the authors and does not necessarily represent the official views of the National Institutes of Health.

## Appendix

### 6 Dale’s backpropagation update

This section studies in detail the Dale’s backpropagation update rule, beginning with the explicit derivation of the same. The following subsections detail the algorithm for implementing this update in a gradient descent framework, and provide proofs regarding the optimality of the resulting weight matrix projection under the Frobenius norm.

#### 6.1 Derivation

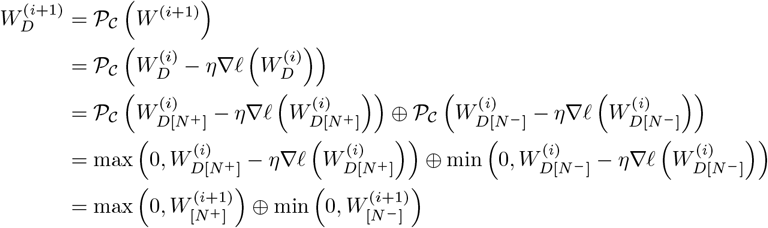

#### 6.2 Algorithm

##### Algorithm 2

Dale’s Backpropagation Update Rule (under the gradient descent optimization scheme)

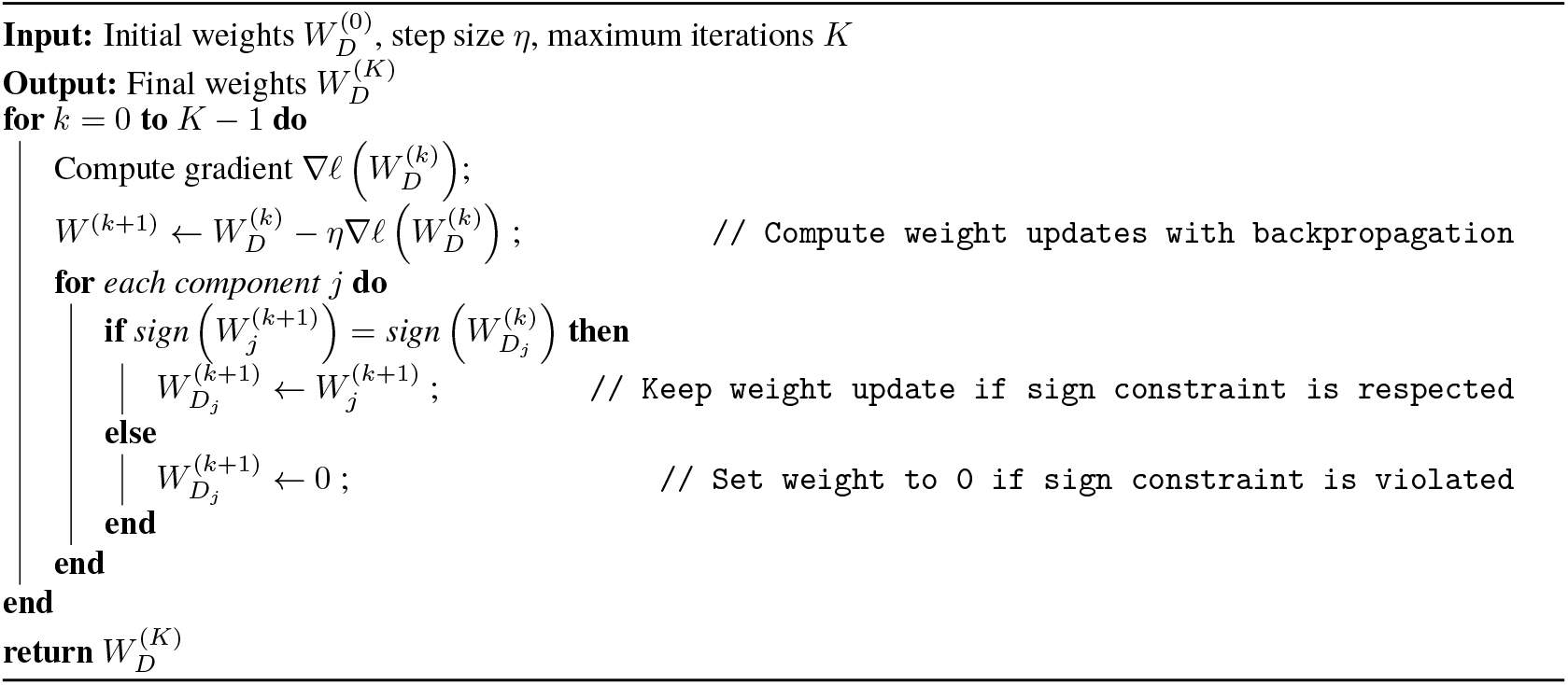

#### 6.3 Closest sign-constrained projection under the Frobenius norm

##### Theorem 5

(Dale’s backpropagation provides the closest sign-constrained projection of *W* under the Frobenius norm). *Let W* ∈ℝ^*N×N*^ *be a real square matrix. Define the set S* ⊂ ℝ^*N×N*^ *as, (i) Columns* 1 *to k: All entries are non-negative* (≥0*), (ii) Columns k* + 1 *to N: All entries are non-positive* (≤0*). Then, the matrix W*_*D*_ *obtained by applying the Dale’s backprop update to W is the closest projection of W onto S under the Frobenius norm. That is*,

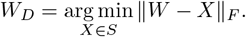

*Proof*. To find the projection of *W* onto *S* under the Frobenius norm, we need to solve:

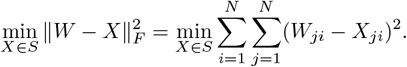

Since the Frobenius norm is separable over the entries of *W* and *X*, we can minimize each (*W*_*ji*_ − *X*_*ij*_)^2^ independently, subject to the sign constraints on *X*_*ji*_:

- **For columns** 1 **to** *k*: The constraint is *X*_*ji*_ ≥ 0.
- **For columns** *k* + 1 **to** *N* : The constraint is *X*_*ji*_ ≤ 0.

**Minimization for each** *X*_*ji*_:

- **If** *j* ≤ *k*:

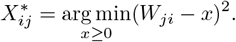

The solution is:

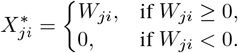

- **If** *j* ≤ *k*:

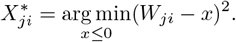

The solution is:

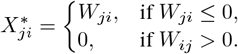

Therefore, the optimal 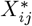 corresponds exactly to the entries of *W*_*D*_ obtained by Dale’s backprop and *W*_*D*_ is the projection of *W* onto *S* under the Frobenius norm.

##### Corollary 6.

*Let W*_*R*_ *be the matrix obtained from W using rectified backprop. Then, the Dale’s backprop matrix W*_*D*_ *is closer to W than W*_*R*_ *is to W under the Frobenius norm:*

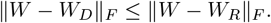

*Proof*. By Theorem 5, *W*_*D*_ is the closest matrix in *S* to *W* under the Frobenius norm. Since *W*_*R*_ ∈ *S*, it follows that:

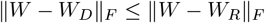

### 7 Alignment of Dale’s backpropagation and Hebbian learning in reinforcing high-magnitude weights

We show that despite the differences in their explicit formulations, both Hebbian learning and Dale’s backpropagation tend to strengthen (i.e., increase the magnitude of) similar weights. In particular, the weights strengthened by Hebbian learning form a subset of those strengthened via the gradient-based Dale’s backprop. Given that weights of higher magnitude inevitably influence the functional connectivity amongst the neurons, preserving weights of higher magnitudes implies preserving those weights which would’ve been important from a statistical learning perspective as well being biologically relevant.

Learning requires a weight *w*_*ji*_ joining pre-synaptic neuron *i* to post-synaptic neuron *j* be changed according to the rule

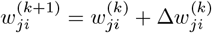

Hebbian learning postulates the update Δ*w*_*ji*_ is given as

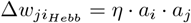

where *a*_*i*_, *a*_*j*_ are the activations of neurons *i, j* respectively while *η* is the learning rate.

On the other hand, the backpropagation update (without loss of generality, in the absence of bias terms) is given as

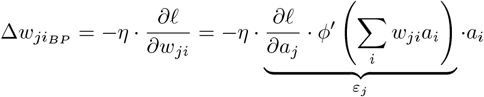

where *ℓ* is the loss function, *ϕ* is the activation function, *ε*_*j*_ is the error corresponding to neuron *j* computed using the chain rule, and *a*_*i*_ has the same meaning as before.

In Dale’s backpropagation, we constrain all activations to be non-negative through a thresholding operation, and weights are restricted to maintain their assigned signs. Under these constraints, the following statements hold true:

**Analysis for** *w*_*ji*_ ≥ 0: In the case of Hebbian learning, given our construction, if *w*_*ji*_ is non-negative, we would need both *a*_*i*_, *a*_*j*_ to be positive to increase |*w*_*ji*_|.

For Dale’s backprop, for a weight *w*_*ji*_ ≥ 0 to increase in magnitude, we require that

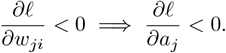

since *ϕ*^*′*^(·) is always non-negative for monotonically-increasing *ϕ* such as ReLU and tanh. This means that as *ℓ* decreases, the neuron *a*_*j*_ contributes positively to reducing the loss. In turn, as learning progresses and reduces the loss *ℓ*, this would lead to an increase in *a*_*j*_ when *a*_*j*_ *>* 0, matching Hebbian learning.

**Analysis for** *w*_*ji*_ ≤ 0 : In the case of Hebbian learning, if *w*_*ji*_ is non-positive, we would require that the *action* of *a*_*i*_ ≤ 0 and *a*_*j*_ ≥ 0 to increase |*w*_*ji*_|.

In the case of Dale’s backprop, for a weight *w*_*ji*_ ≤ 0 to increase in magnitude, we now require

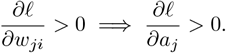

Since 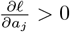 indicates that increasing *a*_*j*_ would increase the loss, the learning process will instead push to decrease *a*_*j*_. Strengthening a negative weight (making it more negative) lowers *z*_*j*_ =Σ_*i*_*w*_*ji*_*a*_*i*_ when *a*_*i*_ *≥* 0, thereby reducing *a*_*j*_ = *ϕ*(*z*_*j*_) and in turn helping to reduce the loss *ℓ*.

This correspondence between Dale’s backprop and Hebbian learning, facilitated by the non-negative activation constraint, suggests that weights strengthened during learning align with those of biological significance. Consequently, when these weights are preferentially retained by our pruning rule, we preserve functionally important connectivity patterns that emerge through biologically plausible learning dynamics.

### 8 Theoretical guarantees for Dale’s backpropagation

#### 8.1 Analyzing convergence of Dale’s backpropagation under the restricted optimum assumption

##### Lemma

(Optimal sign pattern preservation). *Let the vector of learnt weights be W* ∈ℝ^*n*^ *with the components w*_*j*_, *where j* ∈ [1, 2, …, *n*}. *Let L be the Lipschitz constant for the gradients* ∇*ℓ*(*W*), *where ℓ is a loss function. Given a gradient descent-based, component-wise sign-preserving learning rule that uses the projection operator* P_*C*_ : ℝ^*n*^ 1→ ℝ^*n*^ *defined as*

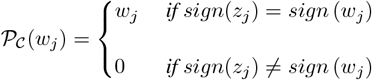

*where* 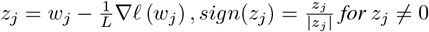, *and sign*(0) = 0. *If sign*(*W*^∗^) = *sign* (*W* ^(0)^) *where W*^∗^ *are the set of weights that can achieve the optimal loss on ℓ, it holds that for any iteration i of regular gradient descent*

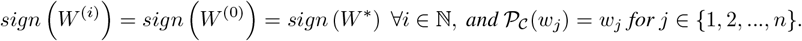

*Proof*. We show by induction that *sign*(*W* ^(*i*)^) = *sign*(*W* ^(0)^) = *sign* (*W*^∗^) ∀*i* ∈ N.

Base case (*i* = 0): The statement trivially holds true since *sign*(*W* ^(0)^) = *sign* (*W*^∗^), by assumption. Inductive hypothesis: For some iteration *i >* 0, *sign W* ^(*i*)^ = *sign*(*W* ^(0)^)= *sign* (*W*^∗^).

To show that for the iteration *i* + 1 it also holds that *sign W* ^(*i*+1)^ = *sign*(*W* ^(0)^) = *sign* (*W*^∗^), we consider *z*_*j*_ as defined, which is the *j*^*th*^ component of

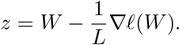

By the Lipschitz continuity of the gradient, we have that

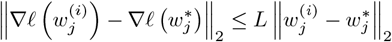

Since *W*^∗^ is the optimal set of weights, we know that 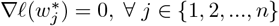. Therefore,

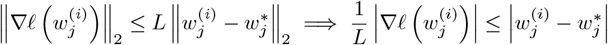

Consider the case where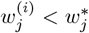.

Here,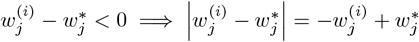. Furthermore, the gradient descent update moves 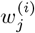 towards 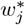 by increasing its value, implying that 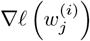 itself is negative. Consequently,

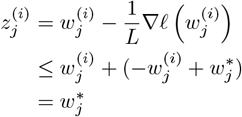

This leads us to the conclusion that 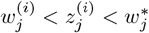.

Since *sign* 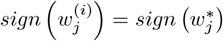 by the induction hypothesis, 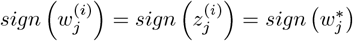 also holds. As a result, 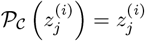 and 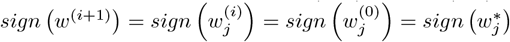.

The case where 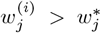 follows similarly, with the difference that since the gradient 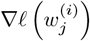 is positive and 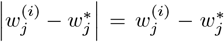, we instead have 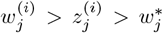. This leads to the same results as before, i.e.,

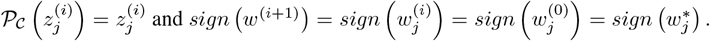

As the choice of the index *j* was arbitrary, these results holds across all indices and therefore

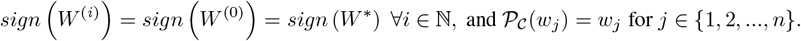

##### Theorem

(Convergence of Dale’s Backpropagation). *Let ℓ be a loss function satisfying the µ*−*Polyak-Łojasiewicz condition, with gradients that are L-Lipschitz such that L* ≥*µ >* 0. *Consider the sequence of weights* 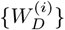 *generated according to the Dale’s backpropagation update, with a step size of* 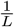. *Given an optimal loss ℓ*^∗^ = *ℓ*(*W*^∗^) = argmin *ℓ*(*W*_*D*_) *where W*^∗^ *has the same sign pattern as all* 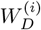 *and a specific error ε >* 0, *it holds for the iteration i that*

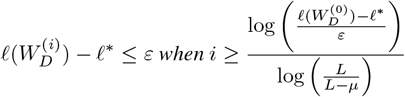

*Proof*. By Lemma 7 we note that the function *g*(*W*) is convex, when it is defined as

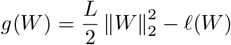

Furthermore, by the first-order equivalence of convexity on *g*(*W*), we have

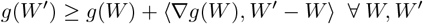

This subsequently implies that

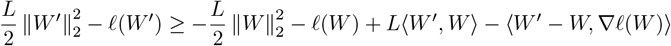

Rearranging terms, we have

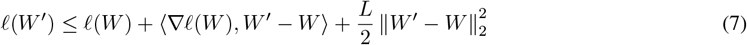

Setting 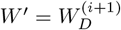 and 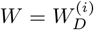 in Eq. 7 while using the Dale’s backprop update rule, we get

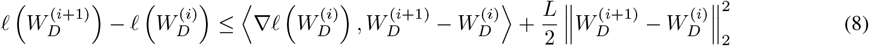

Defining 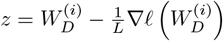 and the projection operator 𝒫_*C*_ as before, Eq. 8 can be re-written as

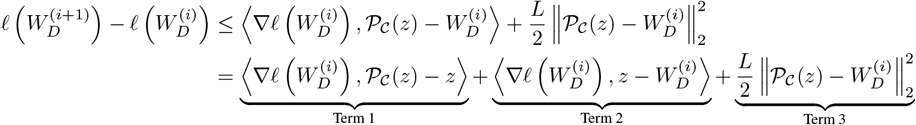

By Lemma 1 we note that **Term 1** is always 0 since 𝒫_*C*_(*z*) = *z*.

Re-substituting 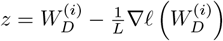 in **Term 2** simplifies it to 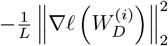

Due to the non-expansive property of metric projections onto convex sets (Theorem 1.2.1 of [98]) and the fact that 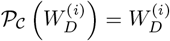 it holds for **Term 3** that

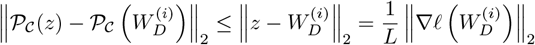

Combining the three terms, we get the bound

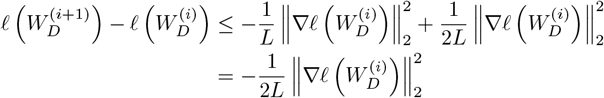

Using the Polyak-Łojasiewicz inequality (Def. 8.1) we get

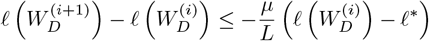

Rearranging and subtracting *ℓ*^∗^ from both sides gives us

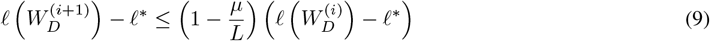

Applying Eq. 9 recursively gives us the result

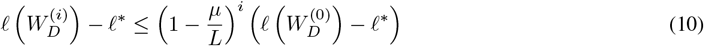

Let the error *ε* be defined as 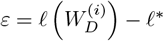 thereby simplifying Eq. 10 to

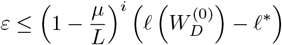

Taking the logarithm on both sides we get

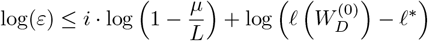

Rearranging the terms finally gives us the bound

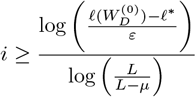

for the target error 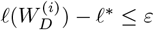.

##### Definition 8.1

(Polyak-Łojasiewicz condition). *A loss function ℓ is said to satisfy the Polyak-Łojasiewicz condition if for some µ >* 0 *it holds that:*

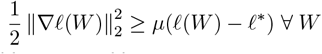

*where ℓ*^∗^ = *argmin ℓ*(*W*) *is the optimal loss attainable*.

##### Lemma 7

(Convexity of transformed function, Lemma 11.1 of [65]). *If the gradient of a loss function ℓ*(*W*) *is L-Lipschitz, then the transformed function g is convex, where g* : ℝ^*n*^ 1→ ℝ *is defined as*

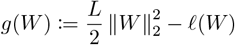

*Proof*. Since ∇*ℓ*(*W*) is *L*-Lipschitz, we have

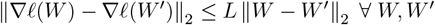

By the Cauchy-Schwarz inequality, we then have

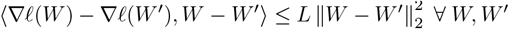

Rearranging terms,

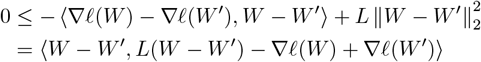

Substituting 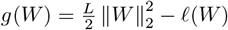 and ∇*g*(*W*) = *LW* − ∇*ℓ*(*W*), we get

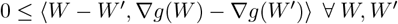

By the monotonicity of the gradient, *g*(*W*) is convex.

#### 8.2 Analyzing Dale’s backprop w.r.t. standard backpropagation

##### Lemma

(Distance between learnt weights). *Let W* ^(*i*)^ *and* 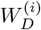 *be the weights at iteration i for standard backpropagation and Dale’s backpropagation, respectively. Assume the gradients* ∇*ℓ*(*W*) *and* ∇*ℓ*(*W*_*D*_) *are upper bounded in magnitude by G and Lipschitz continuous with constant L. Then, the distance between the two sets of weights at any iteration i, denoted as*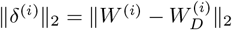, *is bounded by:*

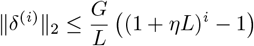

*where η is the learning rate*.

*Proof*. Consider the case where the weights of the network are updated using gradient descent as the optimizer. This implies the following update rules at any iteration *i*:

1. Standard backpropagation update: *W* ^(*i*)^ = *W* ^(*i*−1)^ − *η*∇*ℓ*(*W*)^(*i*−1)^
2. Dale’s backpropagation update: 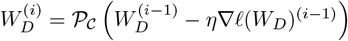

Let 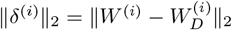 be the distance between the two sets of weights at iteration *i*. We can bound this as:

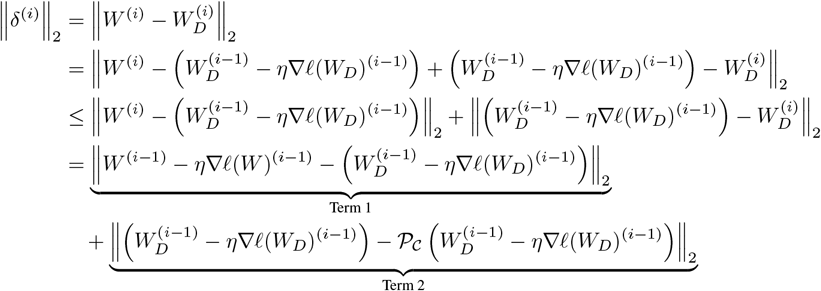

We now bound each term separately.

**Bounding Term 1:**

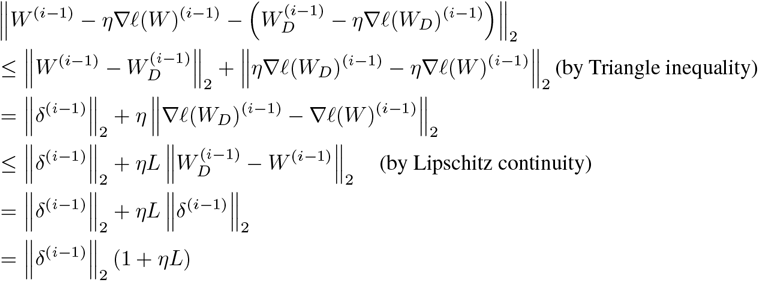

**Bounding Term 2:** The difference between the update using gradient descent before and after the projection step 𝒫_*C*_ at iteration (*i* ™ 1) will never exceed *η ℓ*(*W*_*D*_)^(*i*−1)^ when the update pushes weights *W*_*D*_ outside the feasible region. Therefore,

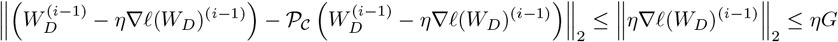

Combining the two bounds, we get:

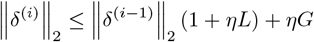

This forms a recurrence relation as follows

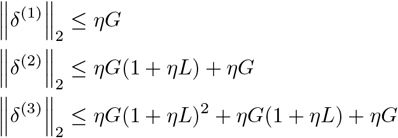

More generally, for any iteration *i*,

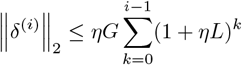

This is a geometric series with ratio (1 + *ηL*) and *i* terms. Summing the series we get

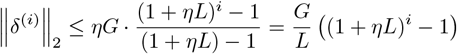

**Theorem** (Differences in errors between solutions). *Let f* (*W*) *be the function represented by a single-layer RNN unrolled over T timesteps, with weights W. Let W*_*D*_ *be the weights learnt using Dale’s backpropagation, and W be the weights learnt using standard backpropagation. Assume the non-linearity ϕ is either tanh or ReLU. Then, the error of the solution found using Dale’s backpropagation with respect to the ground truth y is bounded by:*

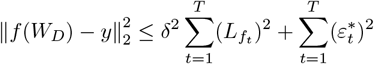

*where* 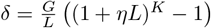 *after K training iterations*,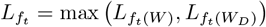 *is the Lipschitz constant of the RNN at timestep t, and* 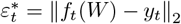 *is the error of the solution found using conventional backpropagation at timestep t*.

*Proof*. We begin by considering a single-layer RNN unrolled over *T* timesteps. Let *f* (*W*) be the function represented by this network, where *W* are the weights. We can express *f* (*W*) as a composition of functions for each timestep:

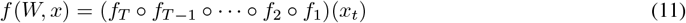

where each *f*_*t*_(*W*_*t*_, *x*_*t*_) = *ϕ*(*W*_*hh*_*h*_*t*−1_ + *W*_*hi*_*x*_*t*_) represents the function at timestep *t*.

Now, let’s consider the Lipschitz constants of these functions. When the non-linearity *ϕ* is either tanh or ReLU (both of which are globally Lipschitz with *L*_*ϕ*_ = 1), it holds by Lemmas 8 and 9 that for every individual layer *f*_*t*_, we have:

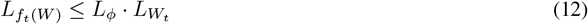

By recursively substituting Eq. 12 in Eq. 11, we notice the following pattern

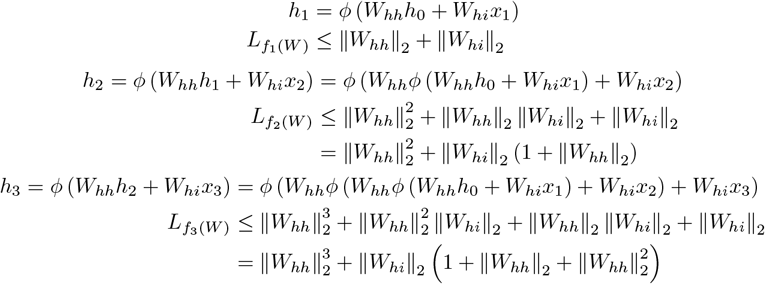

Generalizing the pattern, the Lipschitz constant 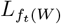 of the RNN at *T* timesteps is bounded by

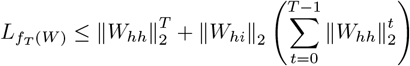

Summing the geometric series we get

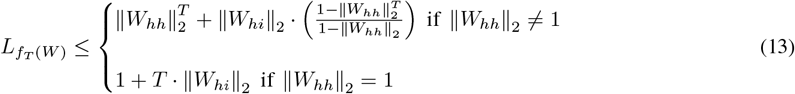

The Lipschitz constant 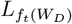 of the RNN with weights *W*_*D*_ can be bounded similarly.

Now, let’s consider the difference between the outputs of the RNNs with weights *W* and *W*_*D*_ at any given timestep *t*. By Lipschitzness,

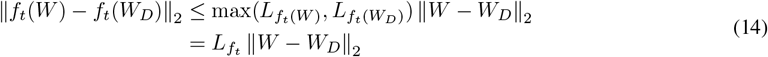

where 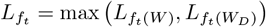.

Applying Lemma 3 after *K* training iterations, we get:

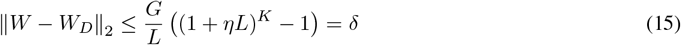

Therefore, we can simplify our bound on the difference between the outputs:

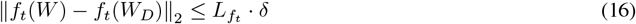

Let *y*_*t*_ be the ground truth at timestep *t*. Applying the triangle inequality, we get:

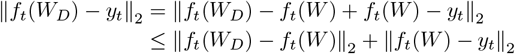

Let 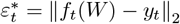 be the error of the solution found using conventional backpropagation at timestep *t*. Hence,

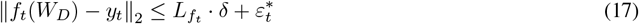

To get the overall error, we sum over all timesteps:

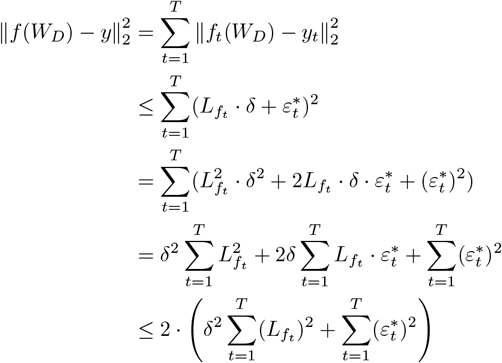

where the last inequality follows from the fact that 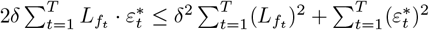 over ℝ.

##### Lemma 8

(Lipschitz constant of matrix multiplication, [99, 100]). *For a linear transformation f* (*x*) = *Wx, the Lipschitz constant L*_*f*_ *is equal to the operator norm of W, i*.*e*., *L*_*f*_ = ||*W* ||_*op*_.

##### Lemma 9

(Lipschitzness of composable Lipschitz functions, [99, 100]). *Let g and h be two composable Lipschitz functions with constants L*_*g*_, *L*_*h*_ *respectively. Then g* ○ *h is also Lipschitz with the constant L*_(*g○h*)_ ≤ *L*_*g*_ · *L*_*h*_.

### 9 Setting *κ* and sparsity values for pruning

#### 9.1 Derivation of *κ*

According to our pruning rule, the probability that a particular edge *w*_*ji*_ is retained in the pruned set is *κ* |*w*_*ji*_|. Noting the fact that all edges in the matrix *W* ∈ℝ^*m×n*^ are sampled independently, the expected number of edges in the pruned matrix *W*^*sparse*^ is simply the sum of probabilities that each individual edge of *W* is retained, i.e.,

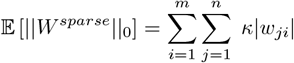

Assuming we wish *W*^*sparse*^ to have a sparsity of *s*, the number of edges in the pruned matrix needs to be (1 − *s*)*mn*, thus giving us the equality

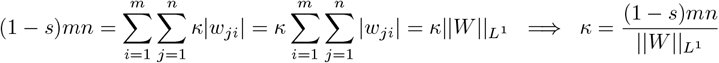

When the matrix *W* represents a recurrent circuit of *N* neurons, 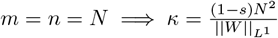.

#### 9.2 Re-normalization of sampling probabilities

Since the initial probability estimates *κ w*_*ij*_ may result in values greater than 1, applying them directly could lead to an overestimation for the probability of retaining certain elements, potentially causing the final sparsity level to deviate from the target. We therefore take a renormalization step to adjust the probabilities so that they sum appropriately, enabling the pruning rule to meet the desired sparsity while maintaining consistency with probabilistic interpretation.

##### Algorithm 3

Probability Re-normalization

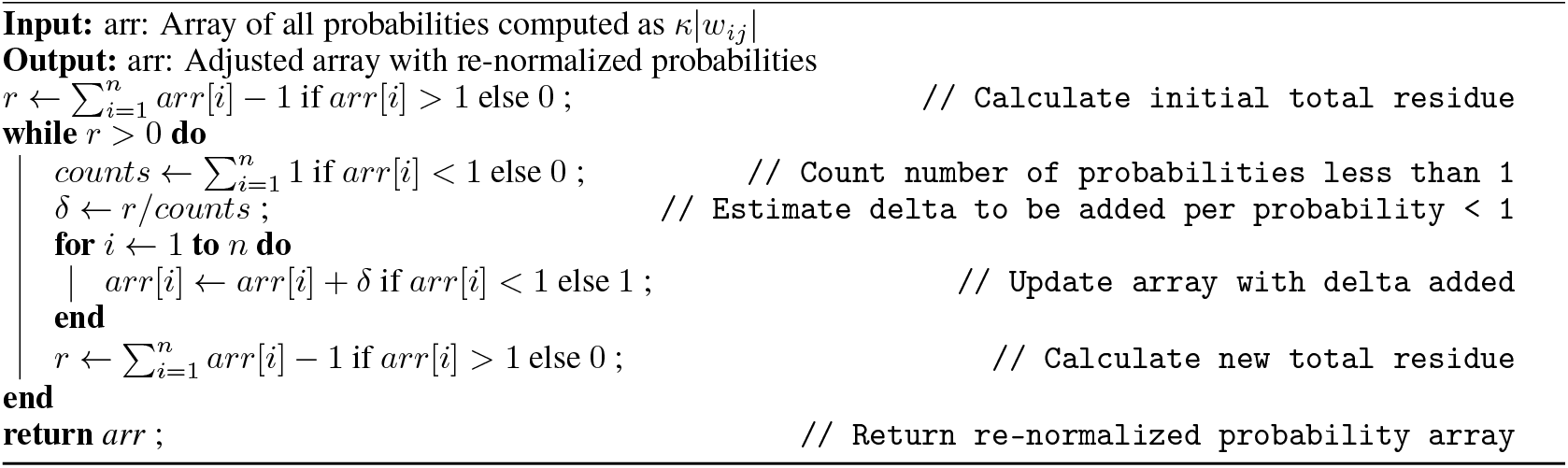

#### 9.3 Adjusting sparsity values

When targeting a specific sparsity level *s* for a matrix (or block of weights), we may find that the matrix *W* already contains a certain number of zero entries, denoted by *z*_0_. If these existing zeros are not accounted for, applying the desired sparsity *s* directly may result in a final sparsity that exceeds the intended level. Therefore, we adjust *s* to a new value *s*^*′*^, which takes into account the current sparsity of *W* as follows:

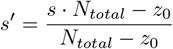

### 10 Expected overlaps with MST across sampling methods

#### 10.1 Lower bounding expected overlap: Top-prob pruned network vs dense MST

Here we establish a (loose) lower bound on the expected overlap between the weights kept when sparsifying an RNN using the top-prob pruning rule and the maximum spanning tree (MST) of the original network, which from the view point of persistent homology, encapsulates all the zeroth-order topological information of a (trained) network.

Consider the square connectivity matrix with *N* pre-synaptic and post-synaptic neurons each. For our purposes we will assume that every neuron always acts as both, a source and a target to at least one other (but not necessarily the same) neuron. The total number of weights in the connectivity matrix is then *N* ^2^ of which (1 − *s*)*N* ^2^ will be sampled for the pruned network to have a target sparsity of *s*. The MST of such a weight matrix will have exactly 2*N* − 1 weights.

Following Kruskal’s algorithm, the probability that the *k*^th^ largest weight by magnitude is in the MST of the bipartite graph can be lower bounded as follows:

In a bipartite graph, the smallest cycle must have at least 4 edges. Therefore, the largest three weights (*k* ≤ 3) are always in the MST.

For *k >* 3 we can lower bound the probability that the *k*^*th*^ largest edge is in the MST. In particular we note that for a weight to lie in the MST, it must not form a cycle, meaning that it either joins two nodes that were both previously not connected to any other nodes in the graph, or joins at most one new node to another connected component on the graph. For the purposes of establishing a lower bound, we only look at the probability of the former.

Consequently, for 3 *< k < N*

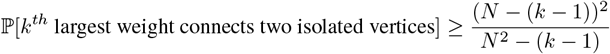

Equivalently,

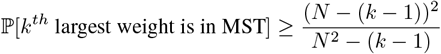

Combining the previous two statements we get the bound:

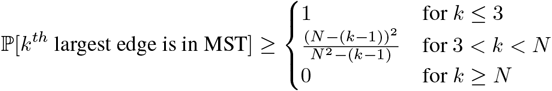

The expected number of weights that overlap with the MST is then simply bounded as

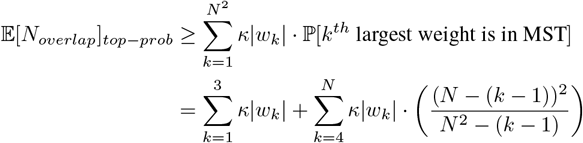

where 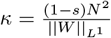

#### 10.2 Expected overlap with MST for random pruning

In the case of random sampling, quantifying the expected number of weights which overlap between the MST and sampled weights is equivalent to that between two arbitrarily chosen subsets, one with 2*N* − 1 weights (i.e., the same size as the MST) and the other with (1 − *s*)*N* ^2^ weights (i.e., the same size as the sparsified connectivity matrix).

To do so we can directly use the expression for the probability mass function of a hypergeometric distribution, where the probability of a random variable *X* having *k* successes (random draws for which the object drawn has a specified feature) in *n* independent draws (without replacement) from a finite population of size *M* objects that contains exactly *K* objects with that feature is given as

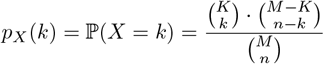

Noting that the MST is a fixed set of 2*N* − 1 weights for any instantiation of the connectivity matrix, the probability that it has exactly *k* weights overlapping with the randomly sampled set of size (1 − *s*)*N* ^2^ out of a possible set of *N*^2^ weights is

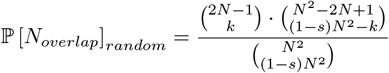

The expected number of sampled weights that overlap with the MST then is simply

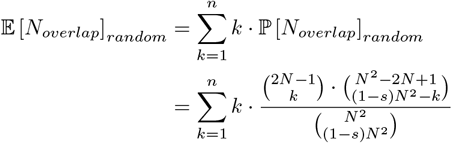

where *n* = min(2*N* − 1, (1 − *s*)*N* ^2^).

### 11 Data pre-processing: Imaging depths

### 12 Connection probabilities within and across cell populations

**Table 1:** Connection probabilities amongst different cell types and populations as per Campagnola et al. [85]. **Columns** = Pre-synaptic population **(source), Rows** = Post-synaptic population **(target)**.

| Connection Probabilities | L2/3<br>Pyr | L2/3<br>SST | L2/3<br>ViP | L4<br>Pyr | L4<br>SST | L4<br>ViP | L5<br>Pyr | L5<br>SST | L5<br>ViP |
| --- | --- | --- | --- | --- | --- | --- | --- | --- | --- |
| L2/3 Pyr | 0.06 | 0.23 | 0.05 | 0.07 | 0.33 | 0.00 | 0.00 | 0.15 | 0.00 |
| L2/3 SST | 0.3 | 0.05 | 0.14 | 0.04 | 0.05 | 0.25 | 0.00 | 0.05 | 0.00 |
| L2/3 ViP | 0.16 | 0.30 | 0.01 | 0.02 | 0.21 | 0.00 | 0.00 | 0.13 | 0.00 |
| L4 Pyr | 0.05 | 0.04 | 0.00 | 0.10 | 0.22 | 0.00 | 0.00 | 0.19 | 0.00 |
| L4 SST | 0.17 | 0.00 | 0.14 | 0.04 | 0.01 | 0.14 | 0.08 | 0.04 | 0.2 |
| L4 ViP | 0.00 | 0.13 | 0.05 | 0.00 | 0.22 | 0.03 | 0.02 | 0.14 | 0.00 |
| L5 Pyr | 0.08 | 0.00 | 0.00 | 0.00 | 0.08 | 0.00 | 0.04 | 0.15 | 0.00 |
| L5 SST | 0.04 | 0.00 | 0.00 | 0.03 | 0.04 | 0.11 | 0.10 | 0.03 | 0.05 |
| L5 ViP | 0.00 | 0.00 | 0.04 | 0.11 | 0.07 | 0.04 | 0.02 | 0.1 | 0.03 |

### 13 Connectivity differences across varying degrees of spatial resolution and types of error

In all three cases we see that the presentation of a novel image (which is a type of expectation violation) increases the inter-areal feedforward connectivity V1 L2/3 → LM L4 in the microcircuit. We see an increase in the feedback connectivity V1 ← LM as well, but the specific feedback pathway that is engaged changes with the type of error; In particular, compared to the Familiar No Change condition, during the Novel No Change condition, an increase in V1 L5 ← LM 5 dominates, while in the case of Familiar Change vs. Novel Change, V1 L2/3 ← LM 5 is the dominant form of increased inter-areal feedback. Across both sets of comparisons, we notice that the novel case leads to increased activity in V1 Vip neurons in both layers 2/3 and 5. In the case of Familiar Omission vs. Novel Omission, we note that there is an increase in the feedback V1 L2/3 ← LM L5 during the Novel Omission condition, as with the other two conditions. However in this case the projection V1 L5 ← LM L5 is lower during the Novel Omission condition, once again speaking to the specificity of feedback projections depending on the type of novelty. As before, we notice that the novel case leads to increased activity in V1 Vip neurons in both layers 2/3 and 5.

In both cases we see that the omission condition increases the inter-areal feedforward connectivity V1 L2/3 → LM L4 as well as both forms of inter-areal V1 ← LM feedback in the microcircuit. Broadly, both no-change vs. omission conditions seem to induce similar connectivty across the various cell-type populations and layers, indicating that the omission, i.e., type of violation of the stimulus, strongly affects the connectivity and subsequent in the microcircuit.

However, at the cell-type level, we see that the strength of interaction from Sst to Vip in L2/3 increases under conditions of novelty and omission, which initially seems at odds with the activity-based findings in [56, 79] that state Vip response is increased during these conditions and Sst activity is reduced. One explanation is that the inferred connectivity is correlative rather than causative, and given that this circuit motif is itself embedded in a larger network with other cell-type populations whose activities we do not have access to in this set of experiments (e.g., PV cells), their absence might skew our results at this finer level when studying inferred connectivity.

## Footnotes

2 During the retraining phase, the synaptic strengths within the network are dynamically adjusted, effectively rescaling the remaining connections to maintain overall network activity and prevent neuron underutilization. This mirrors the biological process of synaptic scaling, ensuring that the network retains its capacity to learn and generalize despite the reduced number of connections.

3 ∥·∥2 always corresponds to the operator norm || · ||op induced by the 2-norm, which is the Euclidean norm if W and WD are vectorized, and σmax(·), i.e., the largest singular value of the matrix if W and WD are considered in their matrix forms.

4 While the methods of [38, 63] using DANNs would allow us to train with sign constraints, unfortunately adapting it to respect structured sparse motifs in non-trivial. We therefore refrain from incorporating any comparisons with such methods in this work.

5 For a more thorough treatment of this topic, we refer the interested reader to Sections 2 & 3 of [73].

6 As well as an overall increase in the connectivity weights targeted to V1 L5 Vip neurons.

7 We note however that these interactions do not directly confirm the experimental results of [56, 79], in that they show a coding change in the relevant populations but not in what would seem to be the same direction, i.e., Sst → Vip reduces in the case of novelty or omission.

